# Multiscale chromatin dynamics and high entropy in plant iPSC ancestors

**DOI:** 10.1101/2023.09.28.559735

**Authors:** Kinga Rutowicz, Joel Lüthi, Reinoud de Groot, René Holtackers, Yauhen Yakimovich, Diana M. Pazmiño, Olivier Gandrillon, Lucas Pelkmans, Célia Baroux

## Abstract

Plant protoplasts constitute the starting material to induce pluripotent cell masses *in vitro* competent for tissue regeneration. Dedifferentiation is associated with large-scale chromatin reorganisation and massive transcriptome reprogramming, characterized by stochastic gene expression. How this cellular variability reflects on chromatin organisation in individual cells and what are the factors influencing chromatin transitions during culturing is largely unknown. High-throughput imaging and a custom, supervised image analysis protocol extracting over 100 chromatin features unravelled a rapid, multiscale dynamics of chromatin patterns which trajectory strongly depends on nutrients availability. Decreased abundance in H1 (linker histones) is hallmark of chromatin transitions. We measured a high heterogeneity of chromatin patterns indicating an intrinsic entropy as hallmark of the initial cultures. We further measured an entropy decline over time, and an antagonistic influence by external and intrinsic factors, such as phytohormones and epigenetic modifiers, respectively. Collectively, our study benchmarks an approach to understand the variability and evolution of chromatin patterns underlying plant cell reprogramming *in vitro*.

## Introduction

Plant tissues have a remarkable plasticity. This phenomenon is illustrated by the capacity of plant parts, tissue fragments or isolated cells *in vitro* to regenerate whole plant individuals. This property is largely exploited by horticulture and agriculture since centuries for the accelerated amplification of garden plants, crop and tree species, the propagation of disease-free plants, the production of plant biomass for industrial applications and the creation of starting material for genetic engineering approaches (Ibanez et al., 2020; Momoko Ikeuchi et al., 2019; Reed & Bargmann, 2021; *Special Issue “New Frontiers in Micropropagation”*, 2021). By contrast, organ regeneration, notably from single cells, is not a prevalent property in the animal kingdom, except in basal lineages like in Cnidarians (Holstein et al., 2003).

Cellular plasticity describes the ability of differentiated cells under certain conditions to reprogramme physiologically and molecularly towards a pluri-competent (or pluripotent) state (Reddy et al., 2021). There is a vivid interest in understanding the mechanisms controlling cellular plasticity. In animals, several pioneer transcription factors have been identified that can potentiate cell reprogramming following overexpression *in vitro* (Iwafuchi-Doi & Zaret, 2014). Similarly, several transcription factors were identified in the model plant Arabidopsis with tissue reprogramming properties (BABYBOOM, WUSCHEL, LEAFY COTYLEDON1, WOUND INDUCIBLE 1 (reviewed in Momoko Ikeuchi et al., 2019; Iwase et al., 2017; Iwase et al., 2011). Among them, so far only LEAFY was formally demonstrated to share the molecular and cellular properties of a pioneer transcription factor (Jin et al., 2021). In addition, several studies concur to the idea that chromatin modifiers, controlling the epigenetic landscape and accessibility, are key to cellular plasticity, in both plants and animals (Birnbaum & Roudier, 2017; Reddy et al., 2021). For instance in Arabidopsis, mutants with reduced levels of DNA methylation or histone methylation (particularly H3K4me4, H3K27me3, and H3K9me2) have altered plasticity and have impaired, or, by contrast enhanced abilities for somatic embryogenesis, callus production, shoot or root regeneration or a combination thereof (He et al., 2012; Ishihara et al., 2019; Jing et al., 2020; Lee & Seo, 2018; Reddy et al., 2021; Shemer et al., 2015).

Conveniently in plants, cells released from aerial, or underground tissues following an enzymatic degradation of the cell wall, protoplasts, provide starting material to generate pluripotent cells. When cultivated on medium supplemented with phytohormones, protoplasts dedifferentiate before reinterring the cell cycle. Proliferation then enables the formation of microcalli, within which some cells will express pluripotency markers. Those plant iPSCs will differentiate shoot and root tissues competent to form a fully fertile plant (reviewed in (Muller-Xing & Xing, 2022); see also Figure 8 in the discussion). Protoplast cultures, considered to share “stem cell-like” properties (Grafi et al., 2011; Sang et al., 2018), and are thus the functional equivalent of iPSC ancestors in plants.

The release of protoplasts from their native tissue rapidly leads to transcriptome reprogramming with a large fraction of affected genes corresponding to stress responses, energy metabolism and photosynthesis (Chupeau et al., 2013). But this response is not uniform, even in cultures composed of 95% of mesophyll cells and the cultures are characterized by a high level of heterogeneity in the transcript composition among cells (Xu et al., 2021). Transcriptome reprogramming is accompanied by profound changes in chromatin accessibility (Xu et al., 2021; Zhao et al., 2001) and in histone acetylation, thought to establish a transcriptionally permissive landscape (Williams et al., 2003). Chromatin changes are also visible at the cytological level. Pioneer studies in tobacco and Arabidopsis (Tessadori et al., 2007; Williams et al., 2003; Zhao et al., 2001) showed that heterochromatin decondenses during, or shortly after protoplasts isolation (Zhao et al., 2001), leading to the spatial dispersion of centromeric and pericentromeric repeats together with their associated DNA and histone methylation marks (Tessadori et al., 2007) and decondensation of rDNA arrays (Ondrej et al., 2010). Despite these large-scale alteration, transposable elements and genomic repeats remain transcriptionally silent, suggesting uncoupling of heterochromatin condensation and silencing (Tessadori et al., 2007). Within the first 3-5 days of culture, heterochromatin gradually recondense and the transcriptome changes again showing more attributes of the cell cycle and regeneration process (Chupeau et al., 2013; Xu et al., 2021).

We aimed here to obtain a comprehensive, quantitative overview of the cytological patterns of chromatin (re)organisation during early culturing of protoplasts corresponding to the dedifferentiation phase (Grafi et al., 2011). In order to foster conceptual comparisons between the cellular reprogramming phases leading to plant and animal iPSC, we will use the term plant iPSC ancestors to refer to the protoplast cultures. Specifically, we deployed high-throughput imaging and customised a supervised-learning image processing to analyse the chromatin patterns in leaf-derived iPSC ancestor within the first 5-7 days of culture. Multivariate analysis of the different chromatin features revealed rapid chromatin changes at different scales. The chromatin of plant iPSC ancestors also rapidly changed in composition, with a notable decrease in linker histone complements. We found that the trajectory of chromatin changes largely depends on nutrient availability and less on phytohormones. In addition, the cultures are characterized by a high heterogeneity as assessed by entropy analyses. Yet, entropy of chromatin patterns decreases progressively in the following 5-to-7 days, a process dampened by the absence of phytohormones but enhanced by an inhibitor of histone deacetylation.

## Results

### H1 as a marker of chromatin reorganisation at early stages of plant iPSC ancestor cultures

To select a live reporter monitoring chromatin changes we analysed various, fluorescently tagged histone variants. First, we considered the three Arabidopsis linker histone variants for which translational fusions are available (Rutowicz et al., 2019; Rutowicz et al., 2015). In a first approach, we scored the number of fluorescent positive cells at day 0 (just after isolation), at day 2 and at day 5. At day 0, H1.1-RFP and H1.2-GFP were detected in most cells (96%, n= 138 and 75%, n= 168, respectively). But at day 5 the fraction of detectable, positive cells had decreased by ∼25% and 22%, respectively (**Figure 1A-B, Fig.S1A**). By contrast, the stress inducible, H1.3-GFP variant reporter was detected only in 2% cells at day 0, likely corresponding to guard cells in which it is constantly expressed (Rutowicz et al., 2015). This fraction did not increase upon culturing (**Figure 1C, Fig.S1A**). Thus, this marker was not further considered. In a second approach, we focused on H1.2 which is the most abundant variant in leaf cells (Kotlinski et al., 2016) and quantified the mean signal intensity per nucleus. We measured a 30% decrease of H1.2-GFP abundance in the fraction of expressing nuclei at day 5 (**Figure 1D**).

**Figure 1.**
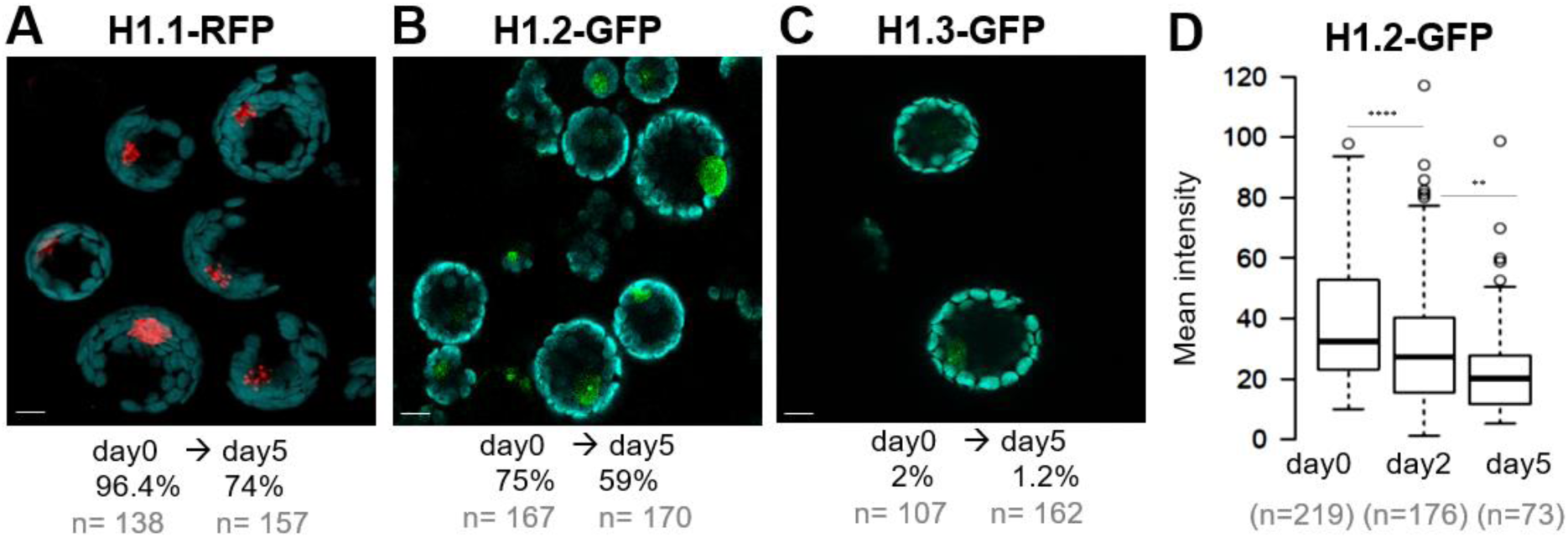
Plant iPSC ancestor cultures are marked by a progressive decrease in H1.1 and H1.2 linker histone variants. Protoplasts were prepared from Arabidopsis leaves expressing a fluorescently tagged variant of H1.1 (A), H1.2 (B) or H1.3 (C) and their level was assessed during five days of culture. (**A-C**) Representative images and percentage (%) of cells with detectable fluorescence signal (n= number of cells scored). See also source data in Table S1 and additional related measurements in Fig.S1. Cyan – chloroplasts (autofluorescence), red – RFP, green – GFP. (**D**) measurements of H1.2-GFP signal intensity in a replicate culture sampled at day 0, 2 and 5. ***, P<0.001; **, P<0.01 (Mann-Whitney-U test). Scale bar 10 µm

In contrast, two core nucleosome histone reporter (H2B-RFP and H3.3-GFP, respectively) were equally detected throughout culturing time (**Fig.S1B**). A GFP tagged H2A.Z reporter also included in the analysis was not further considered since it captured only ∼40% cells (**Fig.S1B**).

Finally, to quantify the mitotic competence of the cultured cells in our conditions, we monitored a S-to-early G2 phase marker (Desvoyes et al., 2020). At day 0 only 5% cells (n=275) showed detectable signal increasing to 8% (n=250) at day 5 and 11% (n=230) at day 6 (**Fig.S1C**). Thus, plant iPSC ancestor cultures are mitotically relatively quiescent in the first 6 days corresponding to the dedifferentiation phase, the first phase of plant cells reprogramming (Grafi et al., 2011). Furthermore, the rarity of S-phase occurrence cannot explain the major decrease in H1 abundance occurring already at day 2. This suggests an active mechanism degrading H1 and likely contributing heterochromatin decondensation described previously in leaf protoplasts (Tessadori et al., 2007).

### A semi-automated pipeline for high-throughput analysis of chromatin reporters

We aimed at high-throughput imaging of plant iPSC ancestors in culture, similar than done previously for analysing cellular morphology (Dawson et al., 2022), but focusing here on chromatin markers. For this, we generated a dual reporter Arabidopsis line expressing both the H1.2-GFP and H2B-RFP markers (selected following the strategy explained above), and used it to establish and benchmark the growth conditions, imaging setup and an image analysis pipeline. The details are provided in the Methods but in short, leaf protoplasts were cultured under sterile conditions in coverglass-bottom 96-well plates for semi-automated imaging using Cell Voyager (**Figure 2A**). The imaging set-up allows to capture up to 60 wells*, i.e.* 60 cultures, per multi-well plate, each being covered by 6 region-of-interest (ROIs). Imaging of a full 60×6 ROIs takes 90 min. After each time point, the plate was returned to the temperature and light-controlled plant growth incubator until the next measurement. Image analysis, illustrated in **Figure 2B**, consisted in a supervised, batch processing approach comprising the following steps performed in TissueMaps (http://tissuemaps.org): (i) maximum intensity projection of the image series; (ii) denoising; (iii) watershed-based segmentation of H2B-RFP-stained nuclei and image masking using the segmented objects; (iv) training, classification and filtering of segmented objects to improve nuclei segmentation; (v) quantitative measurements using validated nuclei objects.

**Figure 2.**
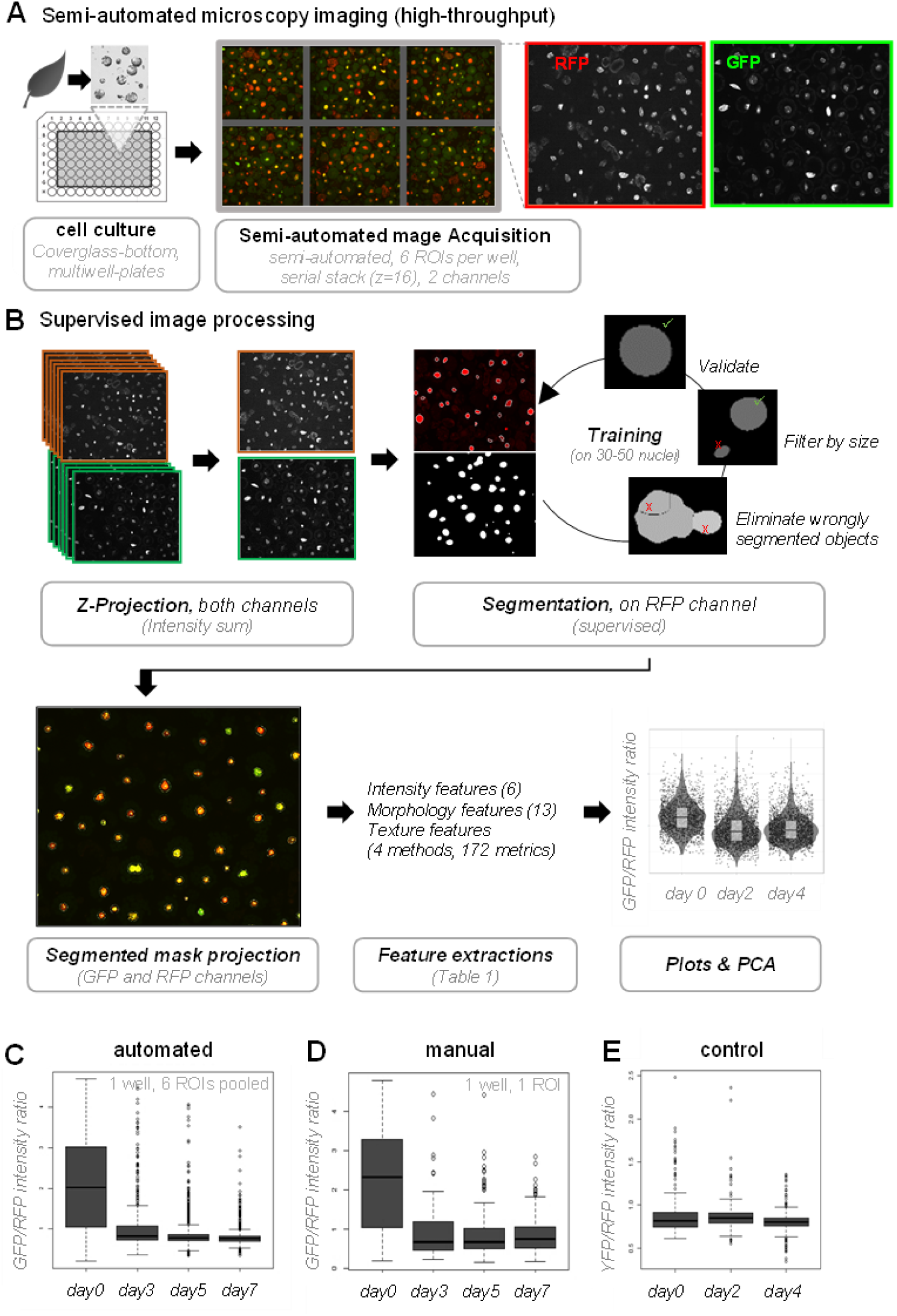
A semi-automated pipeline for high-throughput analysis of chromatin reporters in plant iPSC ancestor cultures. (**A**) Semi-automated microscopy imaging is carried out in cover-glass bottom multi-well plates containing cultures of protoplasts, corresponding to plant iPSC ancestors, using a Cell Voyager platform. Six Regions of Interest (ROI) are randomly selected per well and imaged for each channel. The procedure is repeated for each time point, the culture being returned to the growth incubator in between two measurements (**B**) Supervised image processing followed a workflow as depicted using TissueMaps (www.tissuemaps.org) and enabled the segmentation of several hundred nuclei with high accuracy. The image analysis package returns several descriptors (features) of the segmented objects describing the morphology, the intensity distribution and texture (see details in the text). These metrics can be plotted or further analysed. (**C-D**) Comparison of the H1.2-GFP/H2B-H2B ratios obtained following automated (C) or manual (D) image analysis, showing a reproducible reduction of H1.2-GFP relative to H2B-RFP during culturing time (experiment HTI001). (**E**) Control experiment showing stable signals of a nuclear localised, free YFP marker, relative to H2B-RFP levels (experiment HTI002).

To benchmark this approach, we cultured protoplasts expressing the dual marker in replicate wells and applied our imaging and image processing procedure between day 0 and day 7 (representative images **Fig.S2A-B**). The setup of replicate wells allowed to capture a total of 500-1000 nuclei per time point, per initial culture. Using FDA staining (Saruyama et al., 2013) we confirmed a high density of viable cells at each time points (**Fig.S2C**). However, we observed a decrease in the number of nuclei (cells) identified following the segmentation, particularly at day 7 (**Fig.S2D**), indicating loss of viability, consistently with previous reports (Chupeau et al., 2013; Xu et al., 2021). The analysis of GFP to RFP ratios per nuclei confirmed a dramatic reduction of H1.2 abundance relative to H2B as soon as day 3 (**Figure 2C**). This automated measurement compared very well to a set of manually segmented images used for the same intensity ratio measurement (**Figure 2D**). Then, to assess the reproducibility of the culturing-imaging-image analysis workflow, we set up eight replicate measurements in the same multiwell plate with cultures from two independent reporter lines expressing H1.2-GFP and H2B-RFP. The H1.2-GFP/H2B-RFP intensity ratio distributions were highly consistent between wells at each day (**Fig.S2E**). Finally, as a control, we analysed cultures co-expressing a nuclear localised (nls)-YFP and the same H2B-RFP internal control, using the same setup as before. The analysis showed a stable ratio during culturing (**Figure 2E, Fig.S2F**).

In conclusion, we established a robust automated high-throughput imaging-image analysis workflow enabling the capture of several hundred nuclei per time point suitable for quantitative measurements of chromatin patterns during culturing of plant iPSC ancestor. In addition, the quantifications confirmed that the decrease in H1.2 abundance is a hallmark of chromatin changes within the first two days of culturing.

### Nuclei morphology and chromatin patterns change rapidly

Next, we exploited the numerous image features exported by the pipeline to analyse the cytological organisation of chromatin in plant iPSC ancestor, particularly during the first days of culturing corresponding to the dedifferentiation phase, before cells start dividing (Grafi et al., 2011). Three groups of metrics were produced from the analysis: signal intensity features, morphology features and texture features. This corresponds to a total of 193 metrics per channel and 370 for both channels (morphology features were derived from segmentation on the H2B-RFP signal only, Supplemental Table1). A principal component analysis (PCA, Metsalu & Vilo, 2015) that included all the features indicated clear changes within the first two days of culturing time (**Figure 3A**, replicate **Fig.S3A**).

**Figure 3.**
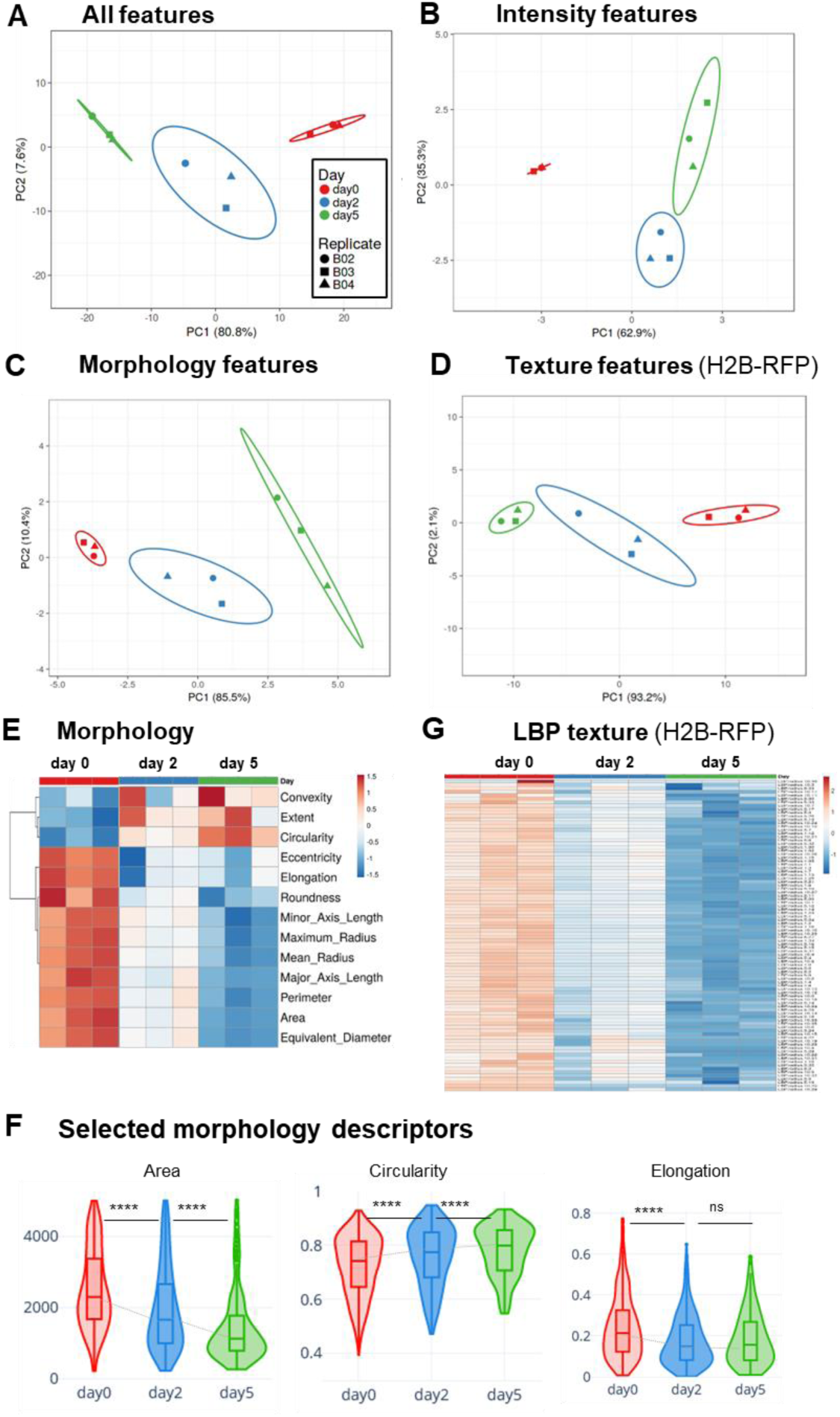
Marked changes in nuclear morphology and chromatin organisation during the dedifferentiation phase. (**A-D**) Principal Component Analysis (PCA) of chromatin features measured in Arabidopsis leaf protoplasts expressing H1.2-GFP and H2B-GFP, imaged ∼4h after release (day0), at day 2 and day 5 (Dataset HTI004). (A) PCA computed on all features (Table S2), (B) PCA on H1.2-GFP and H2B-GFP intensity features, (C) PCA on morphology features, *i.e.* size and shape descriptors of the segmented nuclei, (D) PCA on LBP texture features for H2B-RFP. See also Fig.S3A for replicate PCAs, Fig.S3B for the PCA loading scores per descriptors. X and Y axis show principal component 1 and principal component 2 that explain the given % of the total variance, respectively. Ellipses: 95% confidence interval. Each point represents a culture replicate (well). (**E)** Relative changes for each morphology descriptors during culturing. Heatmaps represent the median value for each descriptor, with unit variance scaling applied to rows. (**F**) Plots of selected morphology descriptors. (**G**) Relative changes for LBP texture descriptors for the H2B-RFP signal distribution. Heatmaps represent the median value for each descriptor, with unit variance scaling applied to rows. Number of nuclei, n= 1008, 508, 238 at day 0, 3, 5, respectively. ****, p<0.0001; ns, not significant, Kruskal Wallis test followed by post-hoc Dunn’s test and Bonferroni correction.

To then estimate the contribution of each feature groups we carried out separate PCA. Clearly, each of the intensity, morphology and texture features contributed to explain chromatin organisation changes during the dedifferentiation phase (**Figure 3B-D**, **Fig.S3A**).

The observation that intensity features distinguish cells at day0, day 2 and day 5 indicates that the relationships between chromatin markers change rapidly in the early culturing phase. Consistent with our previous observation that cells barely divide within the first 5 days of culturing, we found that H2B-RFP intensity distribution did not significantly change during this period of time, but increased, however, at day 7 probably indicating that cells entered S phase (**Fig.S3B**). Likely, the global decrease in linker histone abundance (H1.2-GFP), relative to total chromatin (H2B-RFP), documented before, largely contribute to separate chromatin features on this PCA. Yet, the standard deviation, min and max intensity values also contribute the principal components (**Fig.S3C**) suggesting that, beyond the absolute levels of chromatin markers, their spatial distribution, reflecting the occurrence of chromatin regions with varying density and compaction, also change rapidly.

Morphology features are computed on the segmented H2B-RFP signal, thus provide a proxy for nuclei size and shape (**Fig.S3D**). A PCA considering all morphology features indicated that nuclei undergo continuous size and shape changes between day 0 and day 5 (**Figure 3C, 3E**). Interestingly, although an increase in nuclear size would be expected from the decreased abundance in H1 variants (Rutowicz et al., 2019; She et al., 2013) we did not detect a positive correlation, albeit a moderate one at day 5 (Pearson correlation r <0.3, **Fig.S3E**). Instead, the nuclear size distribution shifts towards smaller sizes at day 2 and day 5 (**Fig.S3F**). The median and the distribution of shape descriptors at day 5 also differ from day 0 (**Figure 3E**), with, for instance, slightly rounder and less elongated nuclei at day 5 than day 0 (**Figure 3F**).

Next, we interrogated the group of texture features, showing a clear evolution during culturing (**Figure 3D**). Textures metrics describe the spatial distribution of signal intensities as a function of scale (Depeursinge et al., 2017a; Di Cataldo & Ficarra, 2017) and can be used to analyse patterns in chromatin organisation (Kerr et al., 2010; Lee et al., 2021; P. Rana et al., 2021). TissueMaps returns metrics corresponding to four types of texture analysis: Gabor wavelet filter, Local Binary Pattern (LBP), Threshold Applied Statistics (TAS), and Hu invariant moment (Hu) (reviewed in (Di Cataldo & Ficarra, 2017), (Hamilton et al., 2007); list Table S2). The LBP analysis, based on neighbouring pixel intensity scanning in incrementally growing circles, is particularly interesting as a proxy of chromatin organisation patterns at different length scales (**Fig.S3G**) (Di Cataldo & Ficarra, 2017; P. Rana et al., 2021). We detected a rapid, global decrease in LBP values along the different radii for the H2B-RFP signal (**Figure 3G**) suggesting a decreasing heterogeneity in chromatin distribution at day 2. The texture metrics of H1.2-GFP signal also showed rapid changes in iPSC ancestor chromatin (**Fig.S3G**). These dynamics in H2B-RFP and H1.2-GFP textures, reflecting the distribution pattern of these histone variants, were confirmed with the TAS and Gabor filter methods (**Fig.S3H-I**). Furthermore, we detected modest, but consistent positive correlations between the LBP metrics and the H1.2-GFP / H2B-RFP ratio, indicating that the changes in chromatin distribution patterns are likely linked with the relative abundance of linker and nucleosome histones (**Fig.S3J**).

Collectively, the analysis of nuclei morphology and of H2B-RFP signal distribution indicate clear changes in nuclear size, shape and in chromatin organisation in the dedifferentiation phase of plant iPSC ancestors (Grafi et al., 2011). These changes occur largely within the first two days and are concurrent, but not correlated, with a decrease in the relative abundance of linker histone H1.2. The distinct texture of H2B-RFP distribution at different length scales at day 5 compared to day 0 suggests a reorganisation of chromatin domains at the (sub)micrometre scale.

### Plant iPSC ancestor cultures show a marked heterogeneity in their chromatin features, which reduces over time

The former analysis clearly showed spread distribution of intensity, size and shape descriptors, indicative of a vast heterogeneity of nuclei type, despite the fact that leaf-derived protoplasts consist in ∼85-90% mesophyll cells (Xu et al., 2021). To assess this heterogeneity, we first generated density distribution maps of chromatin features summarised by the principal components computed previously. The maps confirmed a broad dispersion of the data in the PC landscape, indicating a very heterogenous population in terms of nuclei type and chromatin patterns (**Figure 4A**). The distribution however changed over time with an apparent enrichment of nuclei with similar PC values at day 5 (**Figure 4A and Fig.S4**).

**Figure 4.**
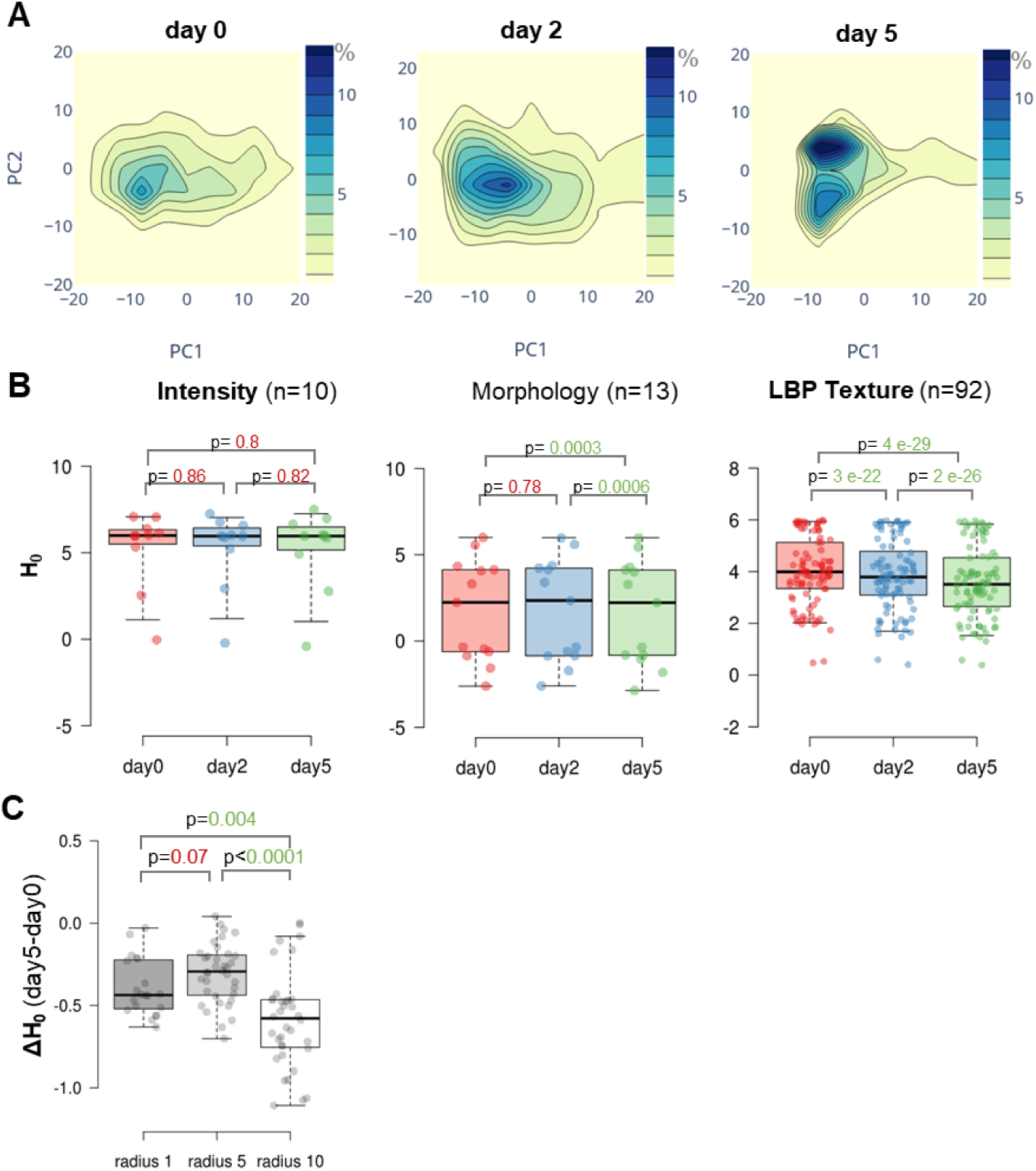
Plant iPSC ancestor cultures are characterized by a high entropy of chromatin features, reducing over time. (**A**) Density distribution of chromatin features summarised by principal components PC1 and PC2 as computed in Figure 3A. Density contours are coloured according to the frequency (percent) of nuclei falling in the corresponding PC space. (**B**) Entropy (H_0_) of chromatin features per family (n, number of descriptors per family). (C) Differential entropy (ΔH_0_) between day 5 and day 0 for LBP texture features of H2B-RFP and at different length scale (radius). *P* values, paired Wilcoxon rank test. Dataset:HTI004.

To quantify this heterogeneity and possible changes during the culturing time, we computed the entropy of the data. Entropy is a useful measure of variability in biological data, capturing both the variance and the shape of the data distribution (Gandrillon et al., 2021). The analysis indeed revealed a marked positive entropy (H_0_) for a large fraction of chromatin features (**Fig.S4B**), changing significantly over time.

We then analysed the family of features separately and found that heterogeneity as assessed by a positive entropy was contributed by all features (**Figure 4B**). Strikingly, the entropy decreased over time, particularly that of texture features and moderately for morphology features, while it remained largely positive for intensity features. Interestingly, the heterogeneity of chromatin distribution, measured by LBP texture features on H2B-RFP, was higher at a small length scale (LBP radius 1, **Fig.S4C**) but entropy reduction over time (ΔH_0_) was more significant for the highest length scale (radius 10, **Figure 4C**).

These findings were confirmed in a replicate experiment and where an additional imaging time point, at day 7 showed that entropy continue to decrease but more slowly after day 5 (**Fig.S4D**).

In conclusion, entropy analysis indicates a profound heterogeneity of nuclei morphology and of chromatin organisation among iPSC ancestors. Strikingly, heterogeneity reduces progressively, mostly within the first five days, and more particularly for chromatin distribution patterns (texture). This suggests a tendency towards homogenisation of chromatin types, although entropy remains high even after 7 days.

### Chromatin heterogeneity is influenced by phytohormones

Protoplast cells cultured in the absence of phytohormones do not grow nor divide and undergo progressive cell death (Zhao et al., 2001). We thus asked whether the chromatin changes detected within the first days of culturing are part of the cellular responses to phytohormones. For this, we partitioned the initial pool of freshly released cells in two media: either in the regular Gamborg B5’s medium (Gamborg et al., 1968) rich in macro– and microelements, vitamins, supplemented with glucose (2%) and phytohormones (auxins and cytokinin); or in the same medium but without phytohormones. First, we asked whether the absence of phytohormones would affect the reduction in the relative abundance of linker histones (H1.2) that was observed previously. Quantifications showed that this is not the case and H1.2 reduction is still taking place in the absence of phytohormones, though perhaps along a milder gradient (**Fig.S5A**). This suggests that H1.2 reduction is not a response to phytohormones in the medium but most likely a response to the cellular isolation, away from the source tissue, and culturing. Next, we interrogated the entire family of chromatin features with or without hormones, using PCA. The analysis showed that cells cultured without hormones undergo similar changes in chromatin organisation than in the presence of hormones (**Figure 5A**, **Fig.S5B**). However, and unexpectedly, we detected that chromatin organisation heterogeneity (measured by the entropy on the LBP texture of H2B-RFP distribution) decreased significantly more in the absence of phytohormones (**Figure 5B**), whereas the heterogeneity of intensity and morphology features were not, or only moderately affected (**Fig.S5C**). We made the same observation when culturing the cells in a nutrient poor medium (W5) without phytohormones (**Fig.S5D**). This suggests that phytohormones contribute maintaining a certain level of heterogeneity during dedifferentiation, in the plant iPSC ancestor cultures. Whether the effect is direct, with chromatin reorganisation responding to phytohormones, or indirect, due to higher cell viability in the presence of phytohormones (Fig.S1D) remains to be determined. Yet, the first scenario is supported by the numerous evidence of crosstalk between phytohormones and chromatin modifiers particularly affecting plant cell identity and plasticity (Maury et al., 2019).

**Figure 5.**
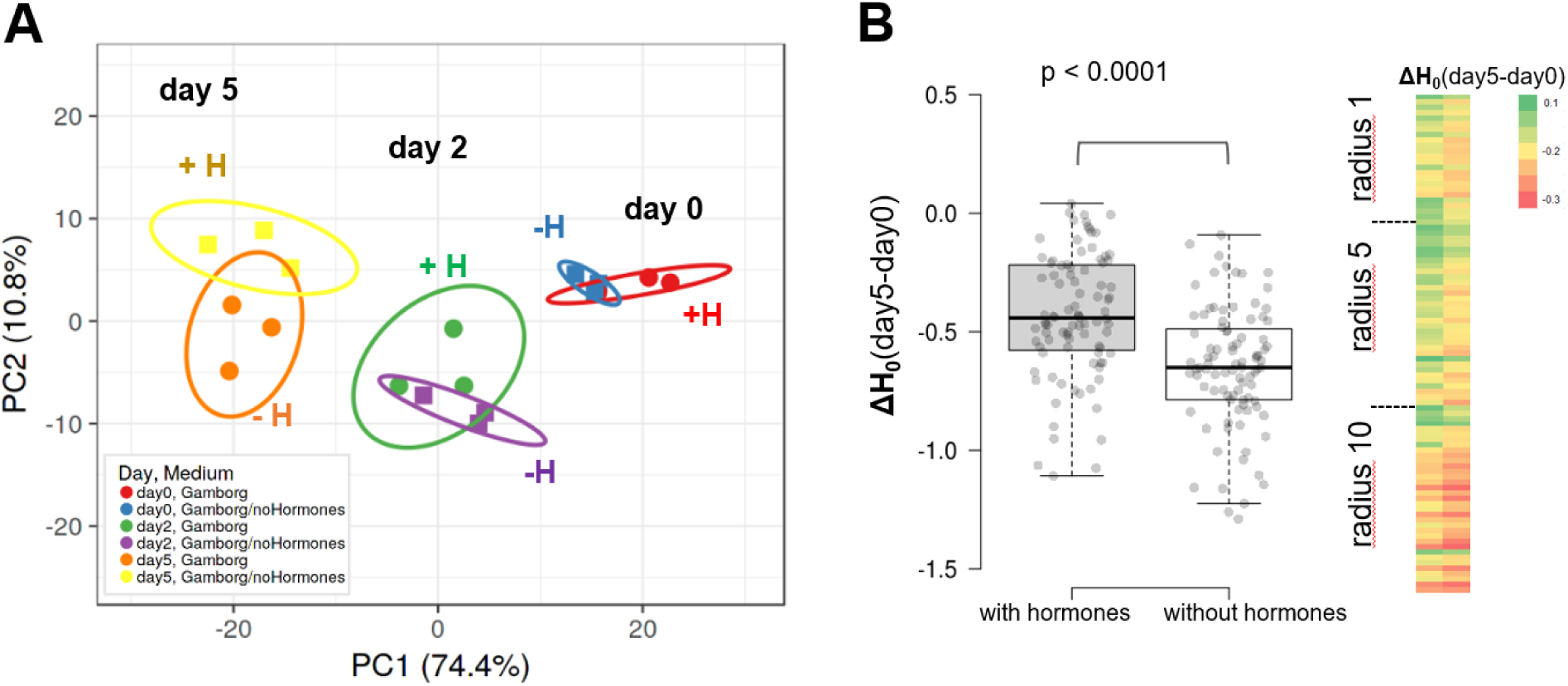
Phytohormones do not influence the trajectory of chromatin changes induced by culturing but dampen entropy reduction. (**A**) PCA of chromatin features of plant iPSC ancestor cultured in the Gamborg medium, including phytohormones (‘Gamborg’ or ‘+H’ as quick annotation on the graph), or in the same basis but without phytohormones (‘Gamborg/no Hormones’ or ‘-H’ as quick annotation on the graph). The PCA was computed with all features (see Methods, Dataset: HTI004). (**B**) Differential entropy (ΔH_0_) between day 5 and day 0 for the chromatin texture features (LBP texture metrics, H2B-RFP) of cells cultured in Gamborg with or without hormones as indicated. P value: Wilcoxon signed-rank test.

### Nutrient availability influence chromatin changes during culturing

To answer the question whether chromatin changes respond to the physiological quality of the culturing medium we partitioned a pool of freshly released leaf protoplasts into W5 (nutrient poor, Yoo et al., 2007) or Gamborg B5 (nutrient rich, (Gamborg et al., 1968), both prepared with or without phytohormones (**Table S3**). We analysed the chromatin features as before focusing on the first two days. Clearly, the culturing medium strongly influenced the chromatin changes with distinct trajectories principally influenced by the nutrient basis more than by the phytohormones (**Figure 6A**, **Fig.S6A**).

**Figure 6.**
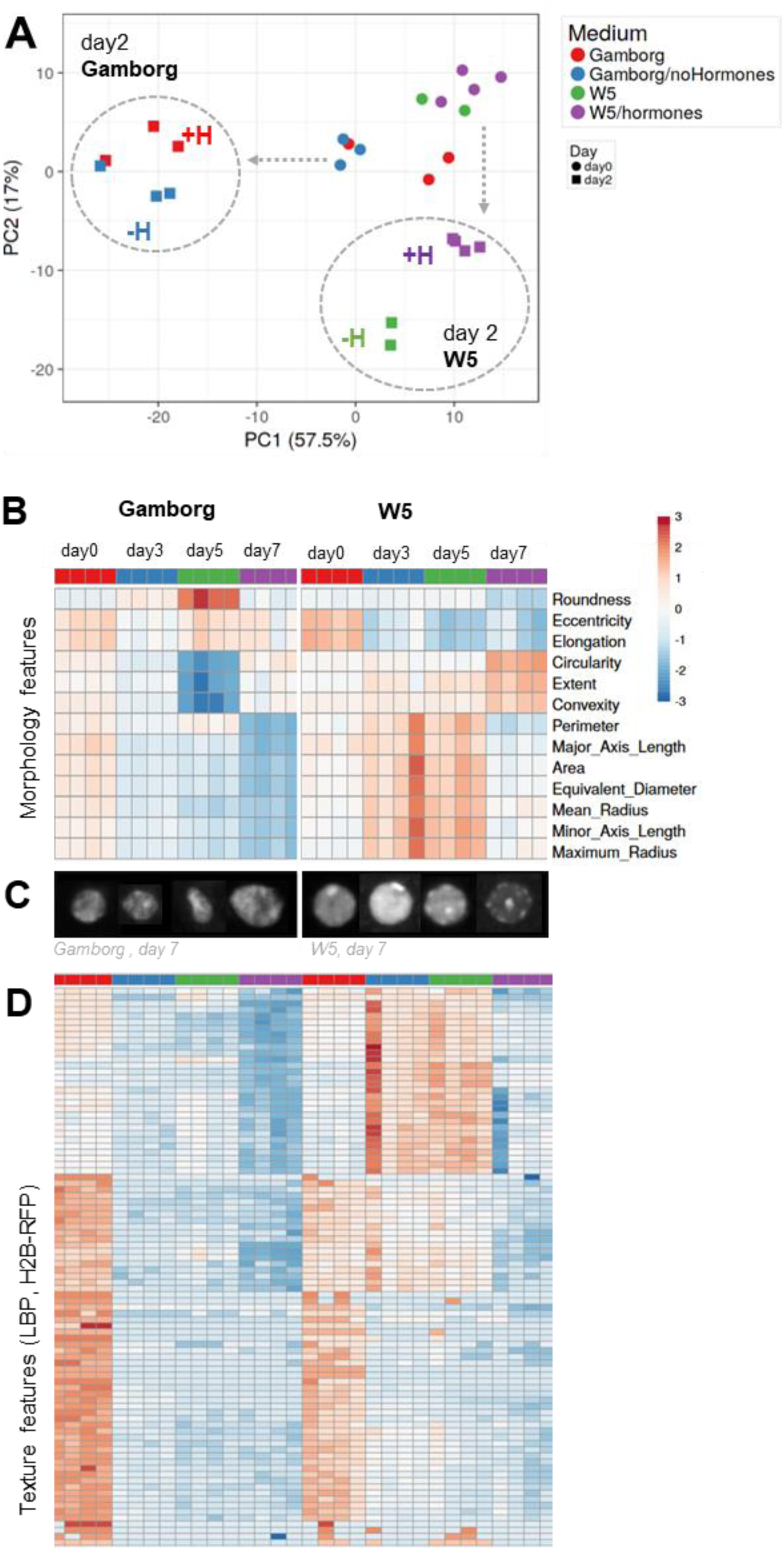
Nutrient availability strongly influences the trajectory of chromatin changes in plant iPSC ancestor cultures. (**A**) PCA of chromatin features in cells cultured either in a nutrient rich (Gamborg) or in a nutrient poor (W5) medium, each with (+H) or without (-H) hormones. The cultures stem from the same, original pool of protoplasts partitioned in the different media and imaged at day 0 and day 2 in two or three replicate wells (number of datapoint with the same colour). Datasets: HTI004 and HTI005. (**B**) Morphological features of nuclei from cultures either in Gamborg or W5 medium, at day 0, 3,5 and 7 (Dataset: HTI001), each square unit is a replicate well. Median values are normalised (centred by rows) and the colour scale shows the fold change. (**C**) Representative nuclei of the same experiment, day 7. (**D**) Chromatin texture features (LBP method, H2B-RFP signal) of the same culture as in B showing comparatively stronger changes between day 0 and day 2 in Gamborg compared to in W5.

Next, to verify whether the nutrient composition influences chromatin changes further in time we imaged new cultures for seven days. The analysis confirmed that major changes occur essentially between day 0 and day 3, yet according to different trajectories depending on the medium, and with chromatin features stabilising rapidly after day 3 (**Fig.S6B**). The typical process of H1.2 reduction also occurred in the nutrient poor medium (W5) although at a slightly lower rate (**Fig.S6C**). The medium affected nuclei morphology (**Figure 6B, Fig.S6D**) with notably rounder and larger nuclei in the nutrient poor (W5) medium (**Figure 6C, Fig.S6D-E**). The nutrient poor medium also dampened changes in chromatin texture across all length scales possibly indicating a slower transition in chromatin reorganisation (**Figure 6E**).

Finally, we asked whether, the heterogeneity of chromatin features may be affected by the culturing medium. The differential entropy (ΔH_0_) between day 0 and day 7 per family of features did not reveal an overall significant effect of the medium (**Fig.S6F**), although, some morphology features showed an increase (*eg* Area) or a decrease (*eg* Roundness) in entropy in the nutrient poor medium compared to the Gamborg’s medium (**Fig.S6G**).

In conclusion, nutrients strongly influenced chromatin dynamics during the dedifferentiation phase of plant iPSC ancestors, with lower nutrient availability inducing rounder, bigger nuclei with a less differentiated chromatin texture.

### Trichostatin A (TSA), an inhibitor of histone deacetylation increases heterogeneity of chromatin patterns

Next, we tested the influence of Trichostatin A, a compound known to inhibit histone deacetylases and increase histone acetylation (Yoshida et al., 1995). TSA treatment was shown to enhance the regenerative competence of lettuce and nicotiana protoplast cultures, notably inducing a higher rate of division starting from day 5 (Choi et al., 2023). We rationalised that this drug could possibly accelerate chromatin decondensation or create changes in chromatin features detectable with our approach. First, a global analysis using PCA did not reveal a major influence of TSA (**Fig.S7A**). Also, TSA did not prevent nor accelerate H1.2 reduction, a typical event of the first culturing day, yet maintained a small fraction of cells with a high H1.2:H2B ratio at day 5 (**Fig.S7B**). The treatment also moderately influenced nuclei size (**Fig.S7C**) but not shape. Possibly, chromatin acetylation as described earlier in tobacco protoplasts (Williams et al., 2003) is so rapid that TSA may act redundantly. However, when we measured the entropy of chromatin features, and more specifically the differential entropy between day 5 and day 0, we observed a strong effect of the treatment Indeed, TSA abolished or strongly diminished the differential entropy for most features (**Figure 7A**), indicating that TSA-treated cultures maintained a high level of heterogeneity at day 5. Hence, histone deacetylation may contribute to channel chromatin reorganisation during the first days of culturing.

**Figure 7.**
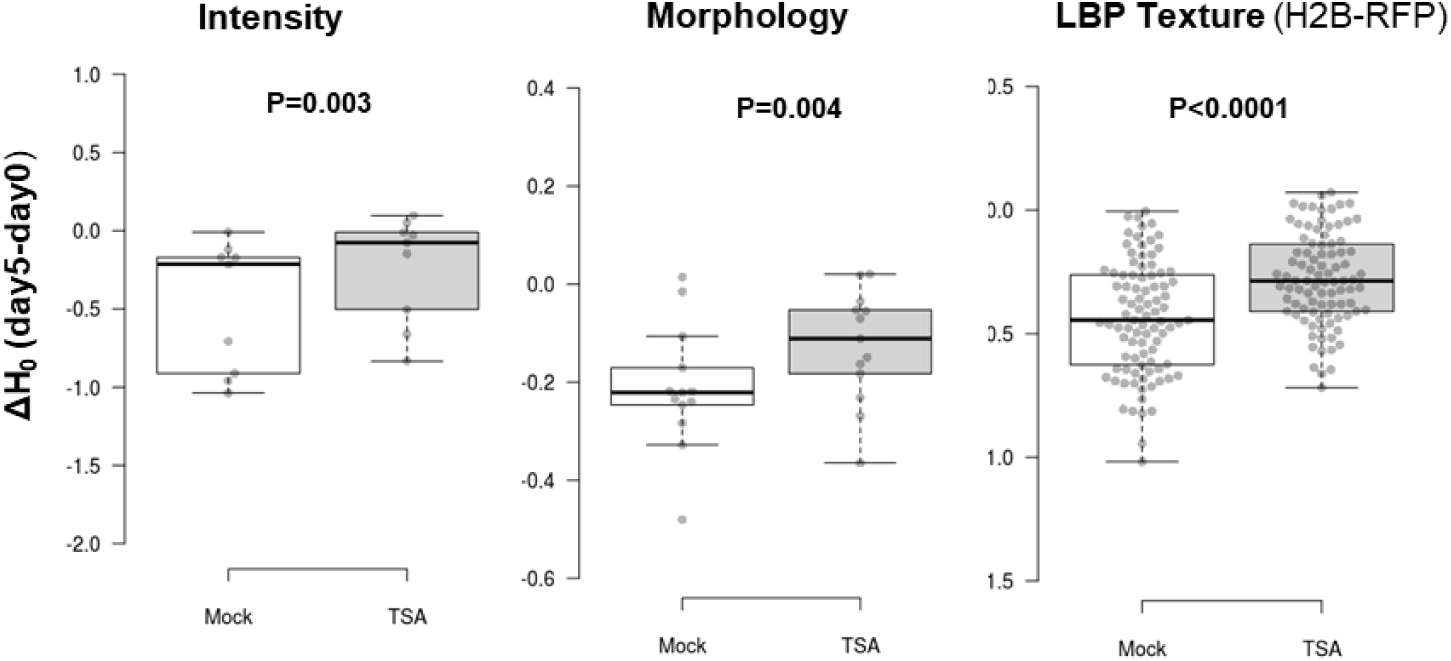
TSA prevents entropy reduction of chromatin features. Differential entropy (ΔH_0_) between day 5 and day 0 for families of chromatin features as indicated, comparing cells cultured in the Mock medium (Gamborg complemented with 2% DMSO) or in the same medium supplemented with 200nM TSA in DMSO. Dataset HTI004. P values: Wilcoxon signed-rank test.

**Figure 8.**
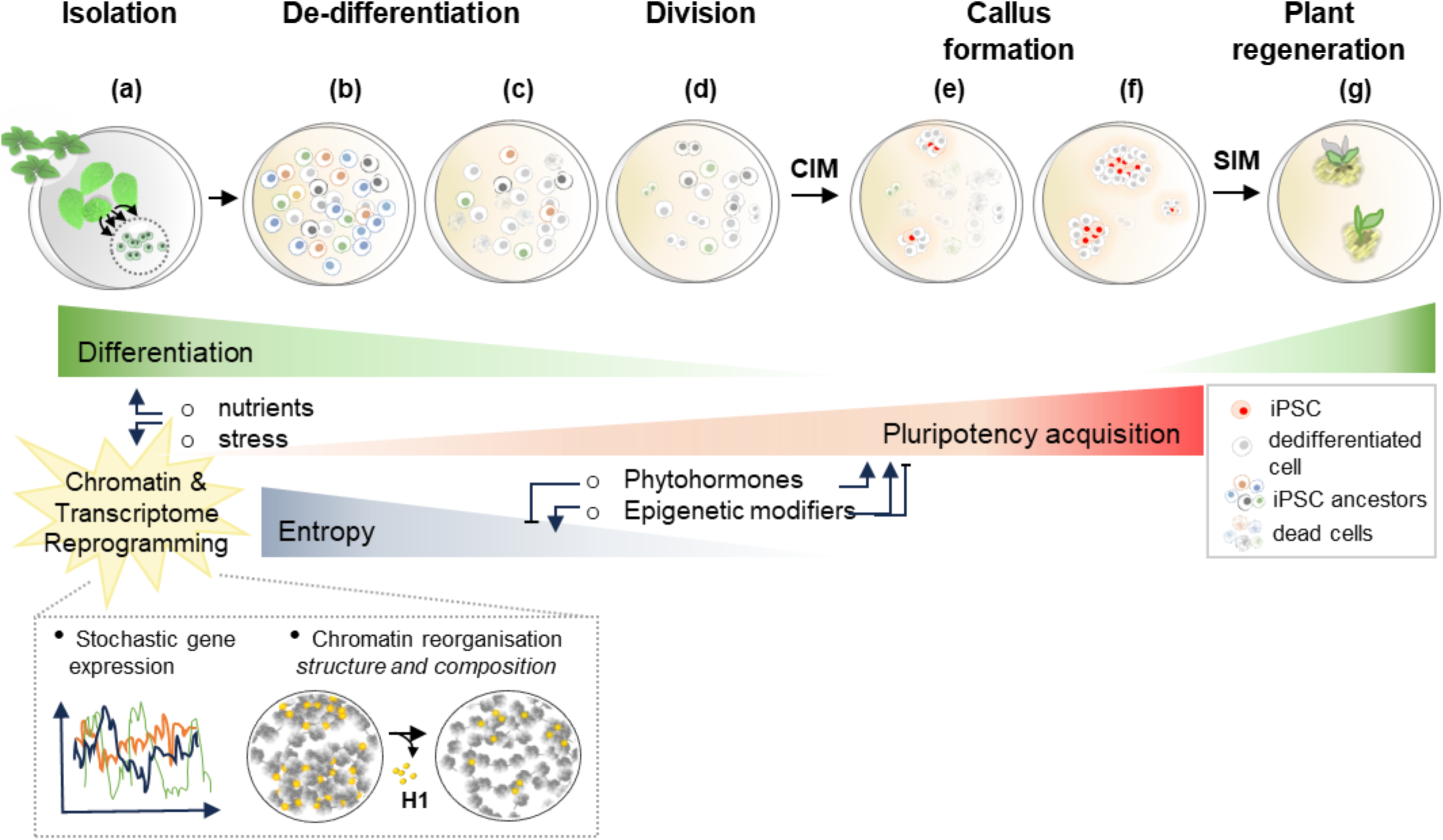
Plant iPSC ancestor cultures are highly heterogenous, with entropy decreasing during early reprogramming. Working model proposing a role for heterogeneity, at the gene expression and chromatin organisation level, in pluripotency acquisition during *in vitro* plant cell reprogramming. This model is proposed based on this work and that of others cited in the main text. Hypothetical extrapolations are made to offer a conceptual framework for future investigations. **(a)** Plant cells devoid of cell walls, called protoplasts, are isolated for instance from shoot tissues (or from other plant organs, not shown here). Protoplasts undergo a phase of dedifferentiation in culture associated with massive transcriptome reprogramming and chromatin reorganisation occurring at multiple scale. Notably, depletion in linker histones (H1) likely impacts structural et epigenetic rearrangement of chromatin domains; H1 depletion affects a large fraction of cells, but not all. **(b)** The initial cell cultures are highly heterogenous (represented with cells of different colours), characterised by a high entropy in chromatin patterns (this work) and stochastic gene expression (Xu et al., 2021), likely induced by the culturing conditions where nutrients and induced stressed contribute. **(c)** In the first culturing days, during the dedifferentiation phase, cellular heterogeneity progressively decreases, and the trajectory depends on nutrient availability, while some cells undergo reprogramming other perish (cells in dashed lines). Entropy (heterogeneity) is influenced by both extrinsic and intrinsic factors, antagonistically: phytohormones have a positive influence on cellular heterogeneity in the culture (the absence of phytohormones accelerate entropy decrease); by contrast, histone deacetylation enables the decrease in cellular heterogeneity, ie may contribute to canalize chromatin (and gene expression) patterns. Whether other epigenetic modifiers act as negative or positive regulators of heterogeneity remains to be investigated. **(d)** After 6-7 days, cells progressively re-enter the cell cycle and (**e-f**) form upon transfer on a callus induction medium (CIM) pluripotent cell masses. Callus cells expressing typical markers of shoot (or root) stem cells, correspond to induced pluripotent stem cells (iPSC) by analogy to animal iPSCs. (**g**) Transfer on a shoot inducing medium (SIM) allow shoot regeneration initiated by the plant iPSC (similarly, roots can be produced from iPSC upon transfer on a root inducing medium, not shown here).

## Discussion

Protoplast cultures offer numerous applications in plant sciences, covering both fundamental and applied research, from the elucidation of molecular and biochemical processes in plant cells to the deployment of new molecular plant breeding approaches, respectively (Xu et al., 2022). They also provide an attractive model to study cellular reprogramming in plant model systems: following release, protoplasts undergo a phase of dedifferentiation (5-7 days) prior re-entering a phase of cell division, which, under an appropriate culturing medium containing phytohormones, can lead to the formation of pluripotent cell masses competent for tissue and plant regeneration (M. Ikeuchi et al., 2019) (Figure 8). Protoplasts, proposed to share a “stem-cell like state” (Grafi et al., 2011) can be considered as *plant iPSC ancestors*, a term conveniently offering a conceptual parallel with animal cells reprogrammed towards iPSC fate. Yet, not all cells are competent for transdifferentiation and regeneration (Pasternak et al., 2020; Sugimoto et al., 2011). In fact, similarly to the low efficiency of animal iPSC production (e.g. 0.01 to 0.1% for human iPSC, Ghaedi & Niklason, 2019), the frequency of cells with regenerative potential was estimated at about 0.5% in an Arabidopsis, leaf-derived protoplast culture (Xu et al., 2022).

In the initial dedifferentiation phase, protoplasts – plant iPSC ancestors – undergo extensive transcriptome reprogramming and large-scale chromatin reorganisation (Chupeau et al., 2013; Moricova et al., 2013; Tessadori et al., 2007; Williams et al., 2003; Xu et al., 2021; Zhao et al., 2001). Surprisingly, even in a relatively homogenous culture composed of 85% leaf mesophyll-derived cells, gene expression changes appear largely stochastic (Xu et al., 2022), raising the question of the level of heterogeneity of cellular states in the plant iPSC ancestor cultures.

We established and validated a semi-automated pipeline for high-throughput quantitative analysis of chromatin reporters at the single cell level, allowing a detailed analysis of the level of heterogeneity of chromatin patterns in plant iPSC ancestor cultures and their dynamic changes during the dedifferentiation phase.

### Harnessing the informative potential of image features provide a new perspective on plant cell’s chromatin organisation

Semi-automated high throughput imaging, followed by supervised image segmentation allowed to record several hundreds of nuclei per day of observation, in 2-4 replicate cultures per experiment. Image analysis computed by Tissue Maps (http://tissuemaps.org/index.html) returned three family of features describing chromatin organisation: Morphology descriptors of nucleus size and shape (the nucleus is the segmented object based on the H2B-RFP signal); Intensity variables for each of the two, jointly expressed, chromatin markers consisting of fluorescently labelled histones H2B and H1.2, respectively; and texture descriptors of the chromatin markers’ distribution.

Morphology and intensity features are commonly used to describe characteristics of segmented cells or nuclei because they are intuitive. Beyond the size, shape descriptors inform here on the roundness, elongation, convexity or solidity of the nuclei, which can be apprehended collectively in a multivariate analysis. Intensity features, in addition, inform on the absolute levels of each chromatin markers (intensity sum), their global compactness or density in the nucleus (intensity mean) and their intensity variability in the segmented object (standard deviation, minima and maxima). By contrast, texture features give abstract representations of signal distribution that cannot intuitively be assigned to a particular distribution pattern, *i.e.* a chromatin phenotype in our case. Yet, those are useful to identify structured and repeated patterns (Depeursinge et al., 2017b; Di Cataldo & Ficarra, 2017). Here, we focused on the LBP method (Local Binary Pattern) that compute a local representation of signal distribution, based on pixel neighbourhood analysis, along circles of incrementally bigger radius, and scanning a growing number of positions along each circle (Ojala et al., 2001). The higher the LBP values at a given radius the more heterogeneous the signal distribution, hence informing on chromatin structures with contrasted density, at different length scales. This approach was reported to be powerful to classify the chromatin types from healthy vs carcinogenic cells presenting different aggregation phenotypes (Priyanka Rana et al., 2021; Venkatachalapathy et al., 2021). Texture analysis indirectly informs on the homogeneity vs aggregation of chromatin at the mesoscale (determined by the LBP method scanning along window of different pixel size) and conceivably also capture the relative proportions of chromatin masses vs interchromatin compartment (Cremer et al., 2020). This approach thus nicely complements former cytological observations describing the disaggregation of the relatively large-scale heterochromatin domains at day 0 of protoplast cultures and their progressive reassembly (Ondřej et al., 2009; Tessadori et al., 2007; Williams et al., 2003; Zhao et al., 2001). We confirmed here that texture features allowed to capture changes in chromatin organisation, at different length scales (see below).

### A rapid, multiscale reorganisation of chromatin which trajectory depends on nutrient availability and less on phytohormones

The release of protoplasts from plant tissues induces dramatic chromatin disorganisation, with the disassembly of heterochromatin domains being a landmark confirmed in different species (Ondřej et al., 2009; Ondrej et al., 2010; Tessadori et al., 2007; Williams et al., 2003; Xu et al., 2021; Zhao et al., 2001). Culturing induces a progressive reassembly of heterochromatin domains and re-entry in the cell cycle (starting day 4 and beyond) in the presence of phytohormones (Tessadori et al., 2007; Williams et al., 2003; Xu et al., 2021; Zhao et al., 2001). Here, high throughput imaging allowed to capture additional characteristics of chromatin dynamics in particularly heterogenous cultures and the role of culturing media. Multivariate analysis identified a clear shift in chromatin features between day 0 and day 2-3, both when considering all family of features (morphology, intensity and textures) or each separately. This indicates that chromatin reorganisation occurs rapidly, early during the dedifferentiation phase, and is detectable at multiple scale. Specifically, changes were most prominent for nuclear morphology, for the intensity level and distribution of H1.2 and H2B and for chromatin texture.

Strikingly, nutrients availability, more than the phytohormones had an influence on the trajectory of chromatin changes. Nutrient mostly affected nuclear morphology and chromatin textures, with H1.2 reduction being largely unaffected. Low nutrient availability prevents protoplasts to re-enter the cell cycle and eventually lead to cell death upon prolonged culturing (Zhao et al., 2001). Here we show that low nutrient availability has an immediate effect and led to larger, rounder nuclei with a less structured chromatin, *i.e.* displaying low textures, even until day 5-7. This is reminiscent of the situation in animal iPSC ancestors where nutrients have a profound impact on chromatin dynamics during cellular reprogramming (Apostolou & Hochedlinger, 2013; Lu et al., 2021). Metabolic fluxes are thought to influence the availability of metabolites used in epigenomic modifications both in plant and animal cells (Apostolou & Hochedlinger, 2013; Lindermayr et al., 2020; Lu et al., 2021; Lu et al., 2023), up to the point where energy metabolism was shown to influence cell fate decisions (Ly et al., 2020). Here, the nutrient rich and nutrient poor media differed not only in the amount of micro– and macronutrients but also in sugar availability. Glucose levels have been shown to control, via the TOR signalling pathway, the cytoplasmic-to-nuclear ratio of *Polycomb*-group repressive complex 2 (PRC2) components influencing H3K27me3 deposition in plants (Ye et al., 2022). H3K27me3 reprogramming is essential for the acquisition of pluripotency in tissue explants primed for callus development *in vitro* (He et al., 2012). Given the ground role of H3K27me3 in cell identity and pluripotency in multicellular organisms, it is tempting to speculate that plant iPSC ancestors may undergo H3K27me3 reprogramming within the first culturing days, corresponding to the dedifferentiation phase (Grafi et al., 2011), under the joint influence of H1 linker histones (Rutowicz et al., 2019; She et al., 2013; Teano et al., 2023) and sugar/nutrient availability (Lu et al., 2023; Ye et al., 2022). Whether other nutrients influence chromatin reorganisation is nevertheless likely given the dramatic impact of nutrient availability on the trajectories of chromatin features.

### A role of linker histone in plant cell dedifferentiation?

The pronounced, decreased abundance of both linker histone variants H1.1 and H1.2 within the first culture days suggests large-scale changes in chromatin composition which may explain the enhanced chromatin accessibility previously measured (Xu et al., 2021; Zhao et al., 2001). Yet, interestingly, while reduced H1 levels correlated with chromatin decompaction and increased nuclear size in differentiated tissues (Rutowicz et al., 2019), this was not the case here, in leaf-derived protoplasts. This suggests mechanisms controlling nuclear size counteracting chromatin decondensation in protoplasts. In Arabidopsis, H1 abundance largely influence the levels and genomic distribution of DNA methylation, the levels of epigenetic marks such as H3K27me3, H3K4me3 and histone acetylation, heterochromatin domains and genome topology (Bourguet et al., 2021; Choi et al., 2020; He et al., 2024; Rutowicz et al., 2019; Rutowicz et al., 2015; Teano et al., 2023; Wierzbicki & Jerzmanowski, 2005; Zemach et al., 2013). Whether the decrease of H1 levels observed here during dedifferentiation *in vitro*, triggers vast epigenomic changes could therefore be expected but remains to be explored.

A link between linker histones, chromatin reprogramming and pluripotency is established in animal cells. On the one hand, gradual and vast changes are observed in the epigenetic landscape, chromatin accessibility and genome topology of animal cells undergoing iPSC reprogramming *in vitro* (Pelham-Webb et al., 2020). Besides, the abundance and type of linker histone variants clearly contribute to several epigenomic and topological features of the genome (Fyodorov et al., 2018). On the other hand, depletion in somatic H1 variants is necessary to *in vivo* pluripotency acquisition in mouse primordial germ cells (Christophorou et al., 2014), and *in vitro* iPSC reprogramming is enhanced when an occyte-specific H1 variant (H1foo) is expressed together with the traditional Oct4, Sox2, Klf4 reprogramming factors (Kunitomi et al., 2016). Thus, it would be interesting to determine in the future whether depletion of canonical linker histone variants plays a role in the reprogramming competence of plant iPSC ancestor cultures and whether non-canonical variants exist that may facilitate, like H1foo, this process. In addition, a comprehensive overview of epigenome reprogramming and topological reorganisation covering the entire process of plant cell dedifferentiation and pluripotency acquisition remains to be established, with single cell resolution to account for the high cellular heterogeneity.

### Chromatin features of plant iPSC ancestors show a high entropy which reduces over time and is antagonistically modulated by phytohormones and histone deacetylation

Transcriptome analyses of protoplast cultures from different plant species, have shown extensive reprogramming compared to their source tissue (Cápal & Ondřej, 2014; Chupeau et al., 2013; Xu et al., 2021). Yet recently, single cell-based reconstructions demonstrated a high heterogeneity of transcriptome patterns even in a relatively homogenous protoplast culture composed of 85% mesophyll cells (Xu et al., 2021). Cell-to-cell variability, particularly in animal system, has been recognized as an important factor controlling the inherent properties of a cellular system prompting investigations to understand its origin and regulation (Eldar & Elowitz, 2010; Eling et al., 2019; Guillemin & Stumpf, 2021; Huang, 2009; Mitchell & Hoffmann, 2018; Mojtahedi et al., 2016; Pelkmans, 2012; Richard & Yvert, 2014; Safdari et al., 2020). One approach to measure the heterogeneity, or information content, of biological systems is based on the calculation of the Shannon entropy borrowed from statistical mechanics (Gandrillon et al., 2021; MacArthur & Lemischka, 2013). It has been successfully applied to measure variability vs robustness of gene expression during cell differentiation (Dussiau et al., 2022; Richard et al., 2016; Stumpf et al., 2017; Wiesner et al., 2018) and reprogramming (Guillemin et al., 2019; Ye et al., 2020). Recently as well, entropy analysis was used to describe heterogeneity of chromatin states at developmentally regulated loci, as an effective information content predicting local genome topology and the competence to binding transcription factors (D’Oliveira Albanus et al., 2021). Here, consistent with stochastic gene expression (Xu et al., 2021), we identified a high entropy of chromatin features among plant iPSC ancestors. This is reminiscent of mouse and human iPSC cultures displaying a high level of transcriptome and epigenetic heterogeneity, thought to correlate with functional heterogeneity (Cahan & Daley, 2013; Yokobayashi et al., 2021).

Interestingly, entropy decreases over culturing time (5-40% depending on features) suggesting a canalization process during this reprogramming phase, towards homogenization, although entropy remains largely positive after 5-7 days culturing. We found that entropy reduction is attenuated in the absence of phytohormones. Although it cannot be excluded that it is a result of higher cell viability in the presence of hormones, it is conceivable that phytohormone-based signalling directly influence chromatin modifiers and remodellers, with an effect on cell identity maintenance and cellular plasticity (Maury et al., 2019).

and promoted when cells are exposed to an inhibitor of histone deacetylation. In contrast, nutrient availability did not influence the heterogeneity of chromatin features. Interestingly, during erythroid cells differentiation stochasticity in gene expression was increased in the presence of a drug inhibiting histone acetylation (Guillemin et al., 2019), rather than deacetylation as in our case. Possibly, the heterogeneity of chromatin patterns and that of transcription profiles are uncoupled to some extent, but more studies on the control of gene expression stochasticity in both animal and plant iPSC ancestor cultures remain necessary.

### Is chromatin entropy as functional determinant of pluripotency acquisition?

A link between chromatin dynamics and chromatin pattern heterogeneity with the actual ability of plant iPSC ancestors to reprogramme remains to be established. Indeed, only a small fraction of cells from a plant iPSC ancestor culture will further develop in a pluripotent cell mass with regenerative ability (Pasternak et al., 2020; Sugimoto et al., 2011). Interestingly, in an Arabidopsis, leaf-derived protoplast culture consisting in 85% mesophyll cells, hence a priori relatively homogenous, only 5% cells seem to contribute an effective regeneration process (Xu et al., 2021). This is reminiscent from the situation in animal iPSC cultures showing variable levels of cellular reprogramming (0.5-10%) depending on the tissue source and inductive method (Kalra et al., 2021; Romanazzo et al., 2020).

Pluripotency in animal iPSC has been proposed to be an emerging property of an intrinsically entropic cellular system, rather than from a unique property at the single cell level (MacArthur & Lemischka, 2013). In this conceptual framework, where pluripotency is a statistical property of a microstate system, uncommitted PSCs undergo weak regulatory constraints leading to a high entropy and stochasticity in gene expression and chromatin states (MacArthur & Lemischka, 2013). As differentiation progresses, more regulatory constraints apply and the heterogeneity of the cellular system diminishes (MacArthur & Lemischka, 2013). Our findings suggest a similar conceptual framework to explain plant iPSC ancestor properties (Figure 8): the release of cell-wall free cells away from the tissue context may abolish regulatory constraints stabilising cellular identity and leading to a high heterogeneity in gene expression and chromatin organisation; appropriate culturing conditions may progressively restore regulatory signals in the culture that may reduce entropy, *ie* canalize gene expression and chromatin organisation patterns (Figure 8). Interestingly in this process, we found that phytohormones, at least the specific ratio and concentration used in our conditions, are dampening this process and nutrient availability does not affect entropy reduction. This suggests intrinsic properties of plant iPSC ancestor cultures to re-establish regulatory constraints reducing the entropy of the cellular system.

In addition, the multiscale heterogeneity of chromatin patterns (as captured by textures) is reminiscent from the finding that variations in local chromatin density underscore the differentiation competence of hESC (human embryonic stem cells, Golkaram et al., 2017). This heterogeneity, which is influenced by genomic contacts but also by DNA free space in the nucleus –a variable intrinsically captured by textures in our cytological analysis – is proposed to reflect variable states of molecular crowding, which in turn controls transcriptional bursts and noise underlying cellular reprogramming (Golkaram et al., 2016; Golkaram et al., 2017).

### Conclusive remarks

Our work opens new perspectives to understand *in vitro* cellular reprogramming and pluripotency in plants. Notably, it is interesting to consider a conceptual framework where cellular variability and the associated chromatin and transcription entropy act as possible driving forces during reprogramming (Figure 8), and where dedifferentiation and pluripotency acquisition result from the property of a cellular system rather than that of single cells as this has been proposed for animal iPSC reprogramming (MacArthur & Lemischka, 2013). Furthermore, whether the regulated abundance, and type of linker histones variants in plant cells also drive chromatin reorganisation and epigenome reprogramming during dedifferentiation and pluripotency acquisition *in vitro* like in animal cells (Figure 8) is an exciting question to investigate.

In a broad sense, protoplasts can be considered the ancestors of plant iPSCs, following reprogramming induced by the culturing conditions, similarly to animal somatic cell cultures that are the ancestors of animal iPSC following reprogramming induced by specific molecular factors. The process of induced cell pluripotency share some common principles in both plant and animal model systems: pluripotency acquisition is largely inefficient (<0.5%) and starting cultures are characterised by a high cellular heterogeneity decreasing over time. This collectively suggest that pluripotency may arise from a population-based, statistical property rather than a single-cell competence (MacArthur & Lemischka, 2013).

## Materials & Methods

### Plant material and growth conditions

The *Arabidopsis thaliana* plant lines expressing fluorescently tagged H1 variants under their native promoters were formerly described (Rutowicz et al., 2019; Rutowicz et al., 2015). To generate the dual chromatin reporter line H1.2-GFP/H2.B-RFP the line *promH1.2::H1.2-GFP* (Rutowicz et al., 2015) was crossed with *promUBQ10::H2B-RFP* (von Wangenheim et al., 2016).

Seeds were surface sterilized and rinsed in sterile water before transferring on the sterile germination medium (0.5 × MS medium, 1% agar). Seeds were placed on the medium with ca 1 cm distance using toothpicks, stratified 2 days at 4°C and grown 3 weeks in long day photoperiod (16 h, 22 °C day/8 h, 18 °C night) and light flux around 100 μM s−1 m−2.

### Protoplast preparation and culture

Protoplasts were isolated from Arabidopsis leaves based on published protocols (Li et al., 2014; Yoo et al., 2007) with some modifications to ensure sterile conditions during isolation and protoplast culturing as described thereafter. The whole procedure was performed under the laminar hood. All solutions were filter sterilized using 0.22-μm filters. Blades, forceps, white pieces of paper, tubes were autoclaved. For pipetting the sterile filter tips were used. After leaf tissue digestion protoplasts were filtered with sterile single-use cell strainers with 70 μm pores. After isolation cells were suspended in intended media (**Table S3**) and distributed into coverglass-bottom, 96-well plates Greiner Bio-One, Ref: 655087 in 100 μl culture per well. The outer wells of the plate were excluded due to the limited field of view and travel range at imaging. Each plate was sealed with 3M tape to avoid drying, wrapped into a layer of aluminium foil and placed in the growth chamber for 5 to 7 days depending on the experiment.

For the Trichostatin A (TSA) treatment, 100 nM or 200 nM TSA in DMSO was added at day 0 in the culture or the equivalent amount of DMSO (2% or 4%, Mock).

To assess cell viability, fluorescein diacetate (FDA) was added at either day 0, day 2 or day 5 before imaging.

### Microscopy imaging

Microscopy images shown in Figure 1 were taken with a laser scanning confocal microscope (Leica SP5, Leica microsystems, Germany). For scoring (graphs Figure 1) the percentage of cells expressing the chromatin markers were scored manually under an epi-fluorescence microscope (Leica DM6000 Leica microsystems, Germany).

For all other figures: leaf protoplasts cultured in coverglass-bottom 96-well plates were imaged using a confocal microscope Cell Voyager (CV7000, Yokogawa), equipped with a 60× water immersion objective (Nikon, NA1.2) using a illumination by 100-200 mW lasers from Coherent (depending on the channel) and filters from Chroma. Images were acquired as 16-bit images using two Neo sCMOS cameras (Andor), pixel size 6.5 µm/M.

For each well, six regions of interest (ROI) corresponding to a field of view of 277 µm x 234 µm with an image format of 2560 x 2160 pixels over 16 z planes with a step of 1 µm. ROIs were randomly chosen without overlap. Imaging time for one well = 90 seconds; Imaging time for one plate (60 wells) = 90 minutes.

### Image analysis

Images were loaded into TissueMAPS (www.tissuemaps.org, code available at https://github.com/pelkmanslab/TissueMAPS) where ROIs were grouped as 2×3 grids per well. Illumination correction based on averaged intensity statistics across all images and maximum intensity projection along the Z dimension was performed in TissueMaps.

Image processing – was done in TissueMaps v0.6.3 for the following steps: (i) gaussian smoothing with a filter-size of 5 pixels; (ii) Otsu-thresholding in a user-defined range using the red (H2B-RFP) channel, followed by binary mask filling and filtering objects < 200 pixels in area to get segmentation; (iii) feature measurements using the measure_morphology, measure_intensity and measure_texture modules in TissueMaps for both channels without the smoothing applied (see Table S2); (iv) classification of mis-segmentations by interactive training of SVMs (Support Vector Machine) in TissueMAPS. For the training, two classes of objects were annotated and created: A, correctly-segmented and B, non-correctly segmented. Then for each class around 40-50 objects were labelled manually. Object morphology and RFP/GFP intensity features were chosen for training the classifier. In particular the following parameters: area, circularity, roundness, elongation, convexity, mean intensity in RFP channel and mean intensity in GFP channel. In total, 100 objects in 10 images (total=1000) were used for training and resulted in 95% segmentation accuracy (n=417), with 4 of the 5% false positive being truncated nuclei at the edge of the image and 1% corresponding to undersegmented nuclei (**Fig.S8**). The description of the SVM algorithm which was used here is available at https://scikit-learn.org/stable/modules/svm.html). Features measurements and object classification were then downloaded from TissueMaps for further analysis. In R (Rproject.org), we further filtered away outlier nuclei with an area >5000 pixels (“giant nuclei”, 1-5% per dataset, possibly from trichomes, Walker et al., 2000).

### Principal Component Analysis (PCA)

PCA were computed using Clustvis (Metsalu & Vilo, 2015). The original dataset exported from Tissue Maps (HTI001, HTI002, HTI004 or HTI005, see Table S4) was subset to remove non informative columns such as redundant identifier codes (related to the plate, experiment and objects), position information (such as Morphology_local centroid_X and _Y, well_position, is_border) and features or experiment description not relevant for the analysis. If too big for upload, the subset data was entered using the input type “paste data”; the data matrix was transposed (“Data matrix Reshape / Transpose Matrix”) and the option “detect column and row annotations” was unchecked to be adjusted manually. Columns with similar annotations were collapsed by taking the median inside each group. Unit variance scaling was applied to rows. SVD with imputation was used to calculate the principal components. X and Y axis show principal component 1 and principal component 2 that explain the given % of the total variance, respectively. Prediction ellipses depict a 95% confidence interval (a new observation from the same group will fall inside the ellipse with probability *p=*0.95).

### Entropy analysis

The initial script for computing Shannon Entropy is described in DOI: 10.1186/s12915-022-01264-9 and is available at https://osf.io/9mcwg/. The adapted script for computing entropy of chromatin features is provided as supplementary information in **SI_File1**. When all cells (segmented nuclei) express the same value for a given feature, this feature’s entropy will be null. The more cell-to-cell variability for a given chromatin feature, the higher value of entropy.

### Plots and statistical tests

Box plots, violin plots, scatter plots, 2D contours and histograms were created using the online tools https://chart-studio.plotly.com/, http://shiny.chemgrid.org/boxplotr/ (Spitzer et al., 2014) or own R scripts (Fig.S1 only). Statistical tests indicated in the figure legends were done using R package or using the online tool https://www.statskingdom.com/mean-tests.html.

## Acknowledgements

We thank Alexis Maizel (COS Heidelberg, Germany) for the Arabidopsis lines expressing the UBQ10::H2B-RFP reporter (von Wangenheim et al., 2016), Alejandro Fonseca for computational help in running entropy analysis, Prof. Ueli Grossniklaus and group members for insightful discussions and Prof Grossniklaus department lab managers and technicians for daily support.

## Funding sources

This work was supported by grants from the Swiss National Science Foundation to CB (310030_185186, IZCOZ0_182949) and to LP (310030_192622), from the European Research Council to LP (ERC-2019-AdG-885579), from the University of Zurich (CB, LP and a postoc grant to KR, K-74502-03-01).

## Data availability

Images used for this study are deposited at DRYAD (doi to be confirmed). The software used for image analysis, Tissue Map, is available at http://tissuemaps.org. The image data used for this study (HTI001-HTI005) are described in Table S4 and available at DOI: 10.5061/dryad.pnvx0k6wp.

## Notes

https://datadryad.org/stash/share/BXm9wlyLx8bupzyXCB6W0WqZ60NMDQvvTcssPSB74jQ.

### Competing Interest Statement

The authors have declared no competing interest.

### Summary of Updates

Semantic change: plant iPSCs was replaced with protoplast or with plant iPSC ancestors when the text aims at fostering comparisons with animal iPSC ancestor cells & cultures Additional explanation to clarify the focus of the study on the dedifferentiation phase and not on pluripotency acquisition Additional Figure 8

