## supplementary figures and tables for "Multiscale chromatin dynamics and high entropy in plant iPSC ancestors"

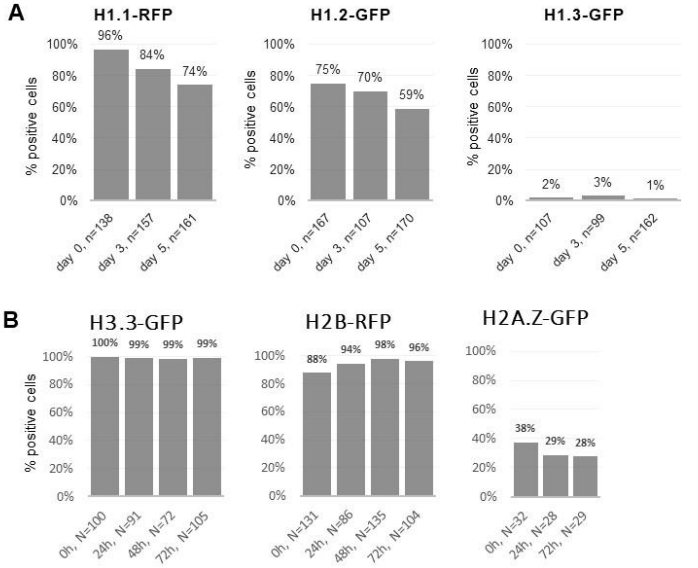


**C % cells expressing the S-G2 cell cycle marker H3.1-mCherry**

| | # positive cells | total (n) | % |
|---|---|---|---|
| **day 0** | 15 | 275 | **5%** |
| **day 1** | 15 | 271 | **6%** |
| **day 2** | 31 | 326 | **10%** |
| **day 5** | 20 | 250 | **8%** |
| **day 6** | 26 | 230 | **11%** |

**Supplemental Figure 1. Expressivity of different histone variants and a cell cycle marker in leaf protoplast cultures**

**(A-B)** Percentage of leaf protoplast showing detectable signal of histone markers (fluorescently tagged histone variants as indicated, see main text and methods for reference and details), at culturing days indicated along the X axis. N= number of cells scored. **(C)** Percentage of cells expressing an S-G2 cell cycle marker H3.1-mCherry (Desvoyes et al., 2020) during protoplast culturing. See also data source in Table S1.

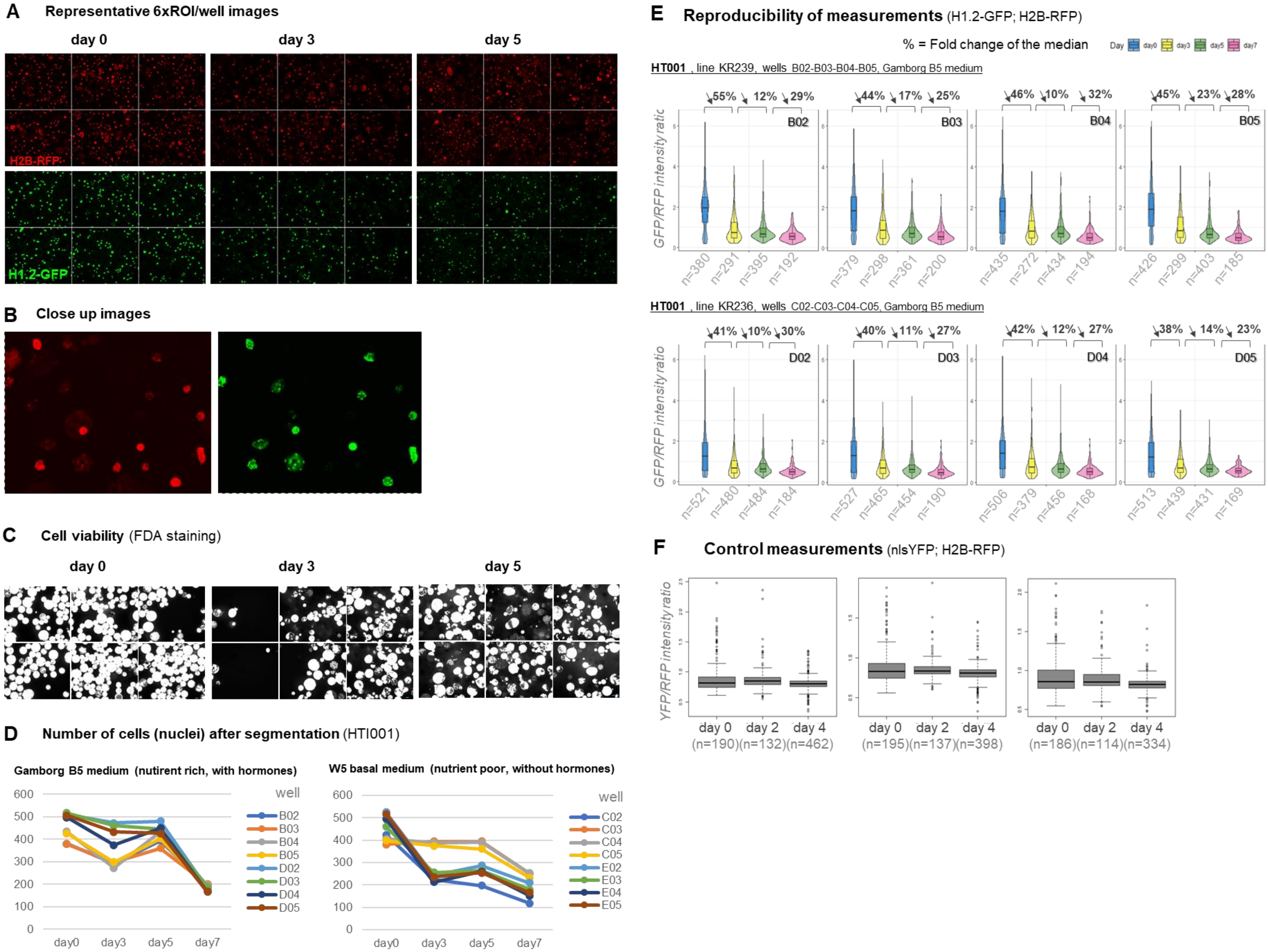
A Representative 6xROI/well images
day 0
day 3
day 5
H2B-RFP
H1.2-GFP
B Close up images
C Cell viability (FDA staining)
day 0
day 3
day 5
D Number of cells (nuclei) after segmentation (HTI001)
Gamborg B5 medium (nutirent rich, with hormones)
W5 basal medium (nutrient poor, without hormones)
well
B02 B03 B04 B05 D02 D03 D04 D05
C02 C03 C04 C05 E02 E03 E04 E05
0 100 200 300 400 500 600
day0 day3 day5 day7
E Reproducibility of measurements (H1.2-GFP; H2B-RFP)
% = Fold change of the median
Day day0 day3 day5 day7
HT001 , line KR239, wells B02-B03-B04-B05, Gamborg B5 medium
GFP/RFP intensity ratio
55% 12% 29% B02 n=380 n=291 n=395 n=192
44% 17% 25% B03 n=379 n=298 n=361 n=200
46% 10% 32% B04 n=435 n=272 n=434 n=194
45% 23% 28% B05 n=426 n=299 n=403 n=185
HT001 , line KR236, wells C02-C03-C04-C05, Gamborg B5 medium
GFP/RFP intensity ratio
41% 10% 30% D02 n=521 n=480 n=484 n=184
40% 11% 27% D03 n=527 n=465 n=454 n=190
42% 12% 27% D04 n=506 n=379 n=456 n=168
38% 14% 23% D05 n=513 n=439 n=431 n=169
F Control measurements (nlsYFP; H2B-RFP)
YFP/RFP intensity ratio
day 0 day 2 day 4
(n=190)(n=132)(n=462)
day 0 day 2 day 4
(n=195)(n=137)(n=398)
day 0 day 2 day 4
(n=186)(n=114)(n=334)

**Supplemental Figure 2. Representative images from leaf protoplasts expressing the dual H1.2-GFP; H2B-RFP markers and cell viability data**

**(A)** Overview of 6 regions of interests (ROIs) per well at day 0, 3 and 5. **(B)** zoom-in images of protoplasts. **(C)** Cell viability assessment at day 0, 3 and 5 with FDA staining. **(D)** Number of cells detected after segmentation showing a reduction towards day 7 of culturing, probably indicating a loss of viability. Comparison of cultures in two different media – the nutrient-rich Gamborg B5 medium and the nutrient-poor W5 basal medium (see Table S3).**(E)** Intensity ratio in protoplasts measured in four replicative wells at day 0, 3, 5 and 7 for two independent transgenic lines expressing H1.2-GFP;H2B-RFP. Arrow: decrease in % in the intensity ratio between two time points. **(F)** Intensity ratio at day 0, 2 and 4 in protoplasts co-expressing the control, nuclear-localised free YFP (nlsYFP) and H2B-RFP. Measurements for three replicates are shown. n, number of protoplasts analysed.

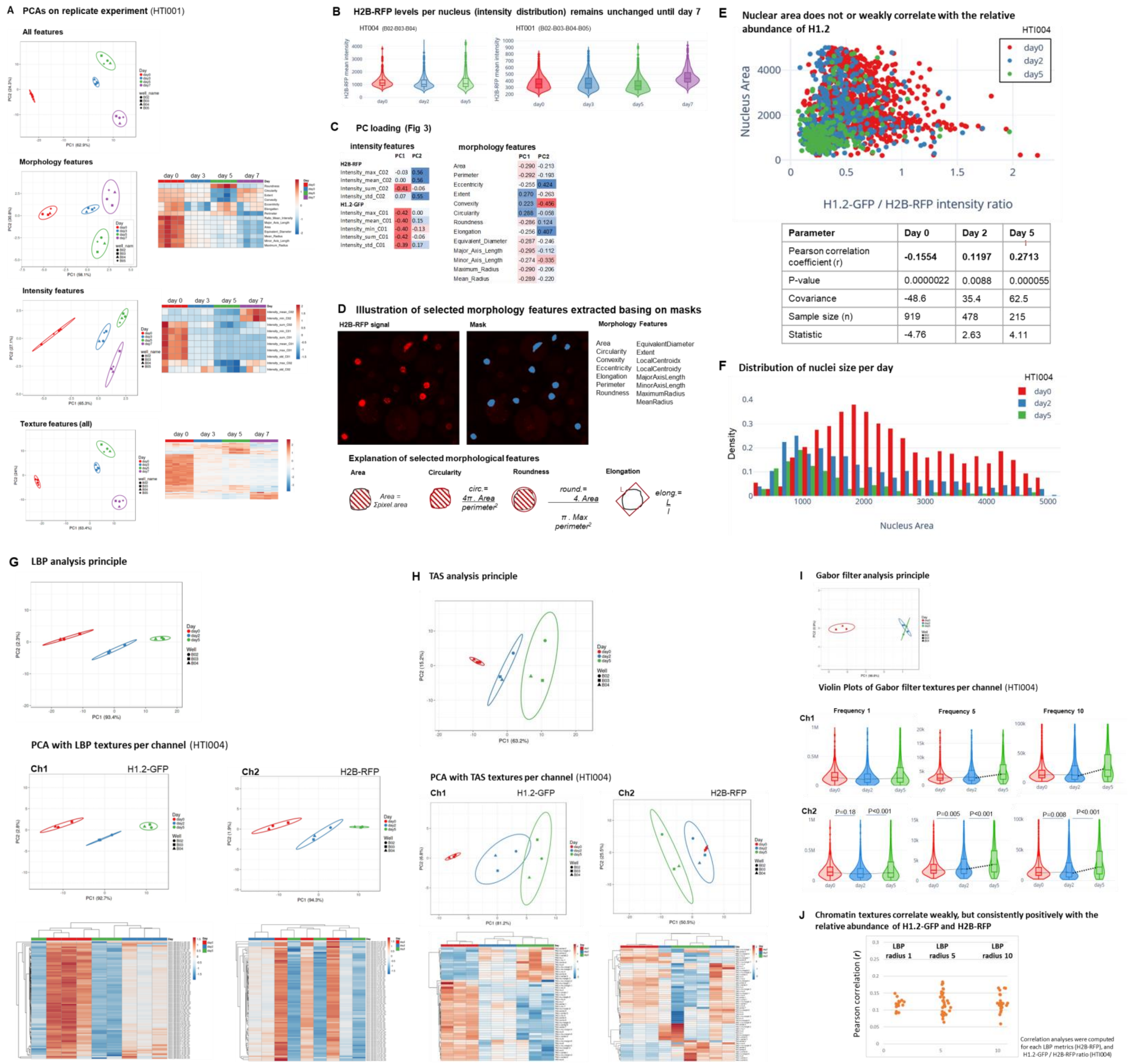


| Parameter | Day 0 | Day 2 | Day 5 |
|---|---|---|---|
| Pearson correlation coefficient (r) | -0.1554 | 0.1197 | 0.2713 |
| P-value | 0.0000022 | 0.0088 | 0.000055 |
| Covariance | -48.6 | 35.4 | 62.5 |
| Sample size (n) | 919 | 478 | 215 |
| Statistic | -4.76 | 2.63 | 4.11 |

**Supplemental Figure 3. Analysis of morphology, intensity and texture features characterizing the protoplast chromatin, in replicate cultures and independent experiments**

**(A)** Principal Component Analysis (PCA) computed on all features, on morphology features only, on intensity features only, or on LBP (Local Binary Pattern) texture features for H2B-RFP only, in protoplasts expressing H1.2-GFP and H2B-RFP at day 0, 3, 5 and 7; dataset HTI001. **(B)** Mean intensity signal for H2B-RFP in protoplasts during culturing in two different experiments HTI001 (sampling at day 0, 3, 5 and 7) and HTI004 (sampling at days 0, 2 and 5). **(C)** List of features contributing to the principal component presented in Figure 3; Blue indicates negative corelations and red indicates positive corelations. **(D)** Selected morphology features extracted from masks, left image – an example image illustrating H2B-RFP marked nuclei, right image – mask obtained based on the left image, bottom panel – illustration of the feature calculations. **(E)** Scatter plot and correlation analysis between nucleus area and H1.2-GFP;H2B-RFP intensity ratio in protoplast. **(F)** Density distribution of nucleus area in protoplast during culturing at day 0, day 2 and day 5. **(G)** PCA computed on the features belonging to the texture class LBP (Local Binary Pattern), bottom panels: PCA separately for each channel. **(H)** PCA computed on the features belonging to the texture class TAS (Threshold Applied Statistics), middle panels: PCA calculated separately for each channel; bottom panel: Heat map representing relative changes for TAS texture descriptors for the H1.2-GFP (left) and H2B-RFP (right) signal distribution. Heatmaps represent the median value for each descriptor, with unit variance scaling applied to rows. **(I)** PCA computed on the features belonging to the texture class Gabor (Gabor wavelet filter), middle and bottom panels showing plots of selective morphology descriptors (Frequencies 1, 5 and 10) for each channel separately; p-values calculated with Kruskal Wallis test followed by post-hoc Dunn's test and Bonferroni correction. **(J)** Pearson correlation analyses computed for selected LBP metrics (H2B-RFP), and H1.2-GFP / H2B-RFP ratio (data set: HTI004).

Additional information for PCA analyses: X and Y axis show principal component 1 and principal component 2 that explain the given % of the total variance, respectively. Ellipses: 95% confidence interval. Each point represent a culture replicate (well). Heatmaps represent the median value for each descriptor, with unit variance scaling applied to rows; Ch1 – channel detecting H1.2-GFP; Ch2 – channel detecting H2B-RFP; See list of features in Table S2.

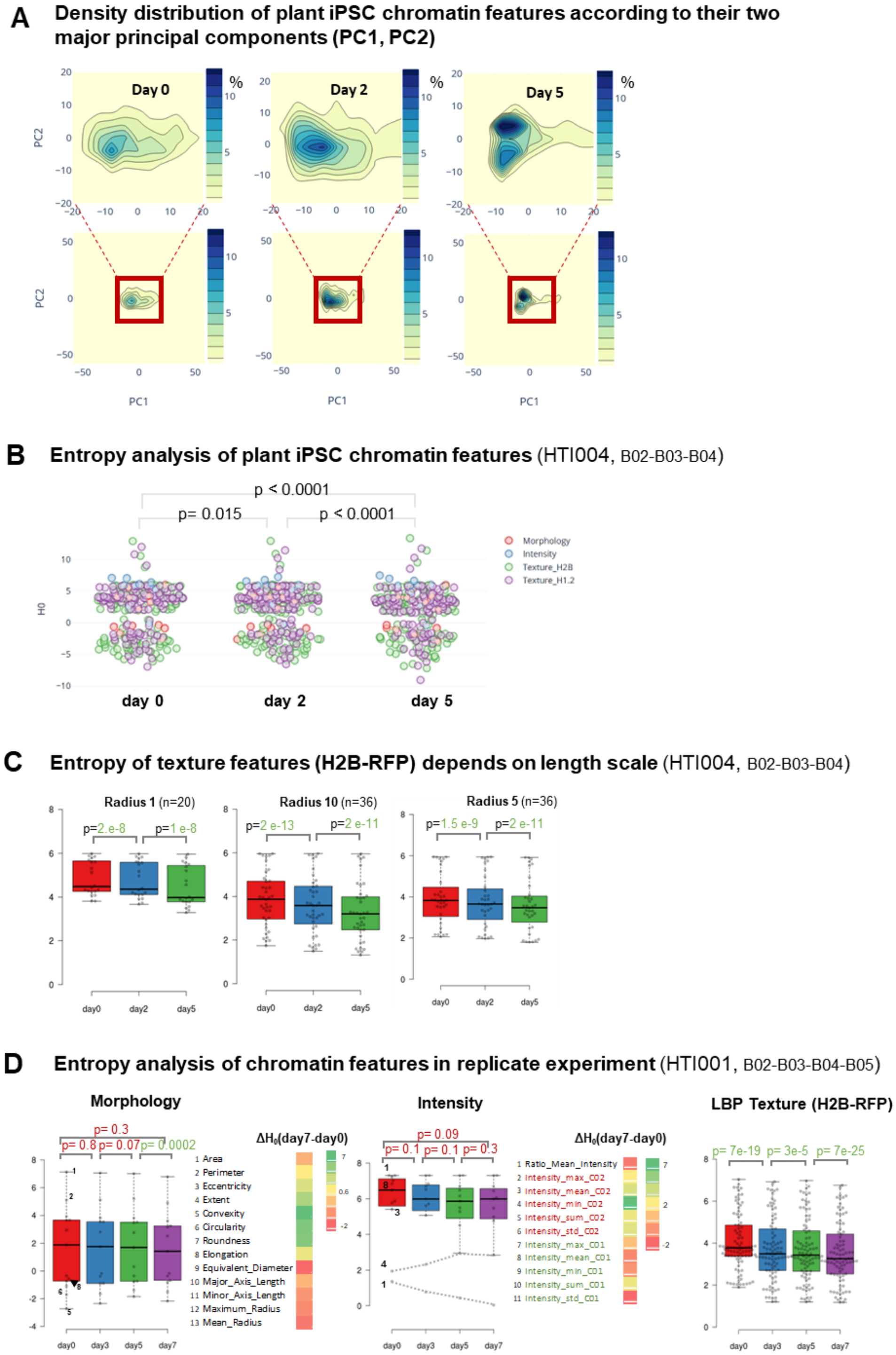


**Supplemental Figure 4**

**(A)** Density distribution of plant protoplast chromatin features according to their two major principal components (PC1, PC2). Density contours are coloured according to the frequency (percent) of nuclei falling in the corresponding PC space. **(B)** Entropy analysis of plant protoplast chromatin features during culturing; dataset HTI004; wells B02-B03-B04. **(C)** Entropy ($H_0$) of texture features (H2B-RFP) for different length scales: Radius 1, 10 and 5, respectively; n, number of descriptors per family; dataset: HTI004; wells: B02-B03-B04. **(D)** Entropy ($H_0$) of chromatin features per family; n, number of descriptors per family; Tukey whiskers extend to data points that are less than 1.5 x IQR away from 1st/3rd quartile. Analysed (those for which the distribution did not allow partitioning for entropy calculation were excluded; $\Delta H_0$, differential entropy; dataset HTI001. P-values calculated from paired t-test; n= number of features; see Table S2 for the list of features and main text for more information.

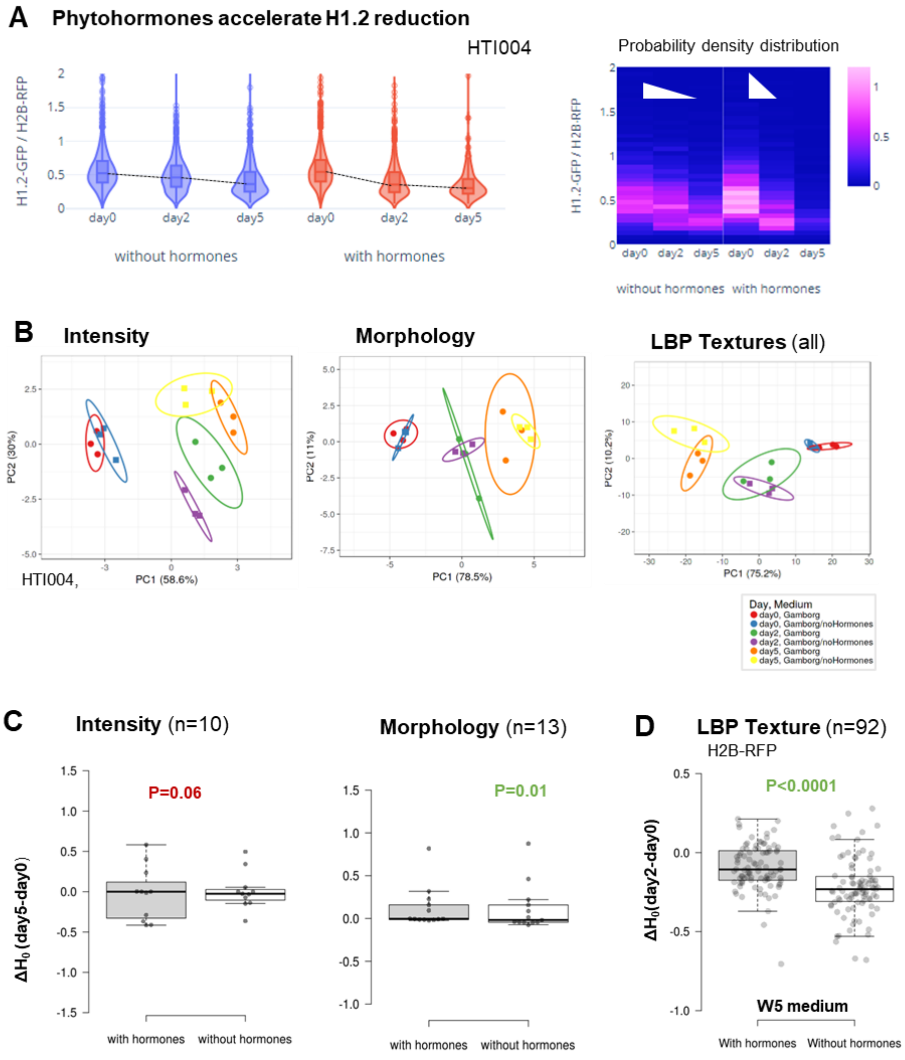


**Supplemental Figure 5. Effect of phytohormones on chromatin features and their entropy in protoplast replicate cultures**

**(A)** H1.2-GFP/H2B-RFP ratio in leaf protoplast cultures in the presence of absence of phytohormones showing a slightly accelerated decreased with phytohormones: dashed lines following the median on the left graph and, heatmap intensity distribution on the right panel **(B)** Principal Component Analysis (PCA) computed on intensity features, morphology features, and LBP texture features for H2B-RFP, respectively during protoplast culturing in the presence or absence of phytohormones. Colours indicate different days and treatment as indicated in the legend. **(C, D)** Differential entropy ($\Delta H_0$) between day 7 and day 0 for intensity and morphology features (C) and LBP textures (D), in leaf protoplast cultured in Gamborg (C) or W5 (D) in the presence or absence of phytohormones. n, number of features per feature group indicated. P value, paired t-test.

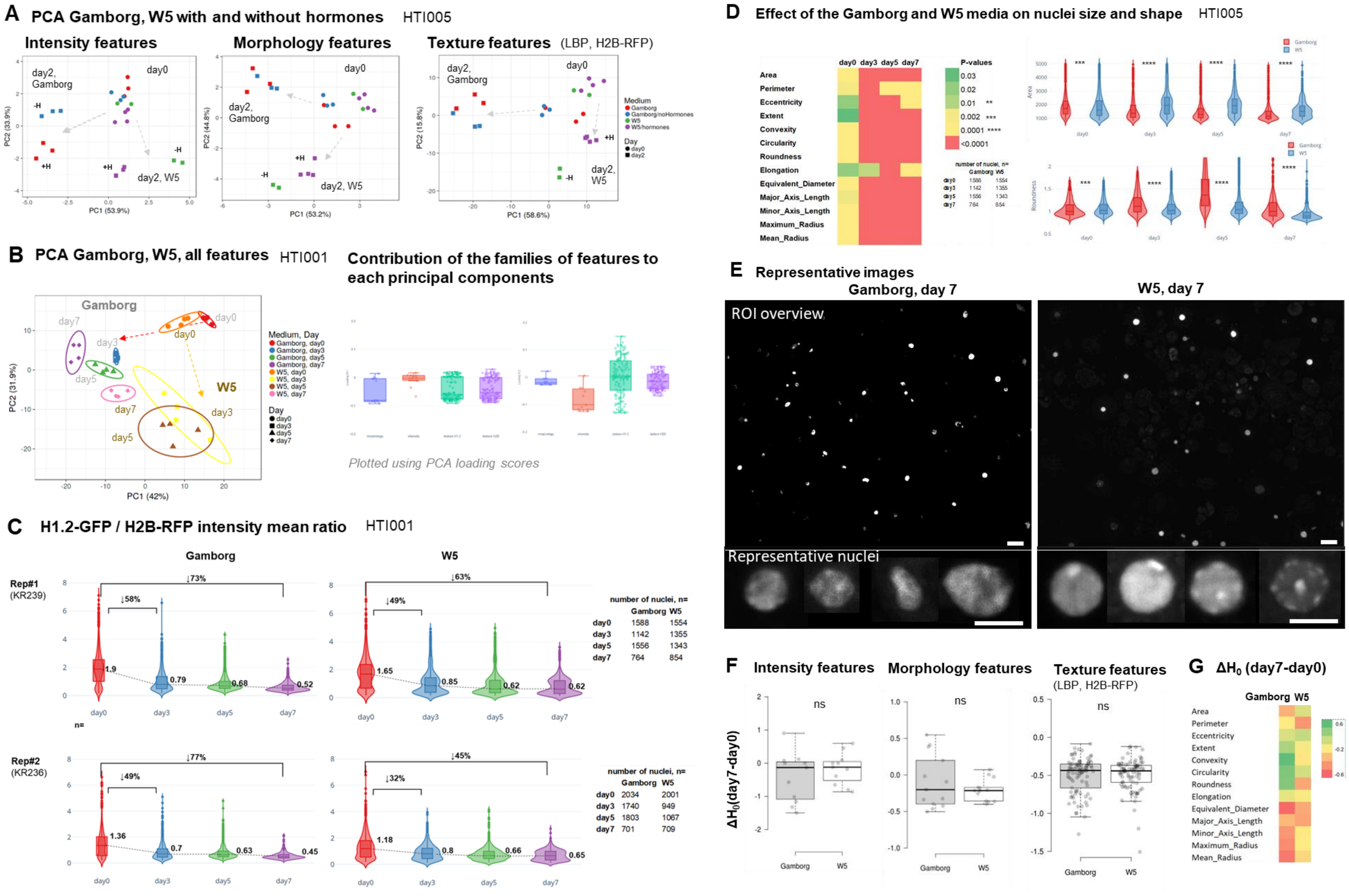
A PCA Gamborg, W5 with and without hormones HTI005
Intensity features
Morphology features
Texture features (LBP, H2B-RFP)
day2, Gamborg
day0
day2, W5
+H
-H
PC1 (53.9%)
PC2 (33.9%)
PC1 (53.2%)
PC2 (44.8%)
PC1 (58.6%)
PC2 (15.8%)
Medium
Gamborg
Gamborg/noHormones
W5
W5/hormones
Day
day0
day2
B PCA Gamborg, W5, all features HTI001
Gamborg
W5
day0
day3
day5
day7
Medium, Day
Gamborg, day0
Gamborg, day3
Gamborg, day5
Gamborg, day7
W5, day0
W5, day3
W5, day5
W5, day7
PC1 (42%)
PC2 (31.9%)
Contribution of the families of features to each principal components
Plotted using PCA loading scores
C H1.2-GFP / H2B-RFP intensity mean ratio HTI001
Gamborg
W5
Rep#1 (KR239)
Rep#2 (KR236)
↓73%
↓58%
↓63%
↓49%
↓77%
↓49%
↓45%
↓32%
1.9
0.79
0.68
0.52
1.65
0.85
0.62
0.62
1.36
0.7
0.63
0.45
1.18
0.8
0.66
0.65
day0
day3
day5
day7
n=
number of nuclei, n=
Gamborg W5
day0 1588 1554
day3 1142 1355
day5 1556 1343
day7 764 854
number of nuclei, n=
Gamborg W5
day0 2034 2001
day3 1740 949
day5 1803 1067
day7 701 709
D Effect of the Gamborg and W5 media on nuclei size and shape HTI005
day0 day3 day5 day7
Area
Perimeter
Eccentricity
Extent
Convexity
Circularity
Roundness
Elongation
Equivalent_Diameter
Major_Axis_Length
Minor_Axis_Length
Maximum_Radius
Mean_Radius
P-values
0.03
0.02
0.01 **
0.002 ***
0.0001 ****
<0.0001
Area
Roundness
E Representative images
Gamborg, day 7
W5, day 7
ROI overview
Representative nuclei
F Intensity features
Morphology features
Texture features (LBP, H2B-RFP)
ns
ΔH0(day7-day0)
Gamborg
W5
G ΔH0 (day7-day0)
Gamborg W5
Area
Perimeter
Eccentricity
Extent
Convexity
Circularity
Roundness
Elongation
Equivalent_Diameter
Major_Axis_Length
Minor_Axis_Length
Maximum_Radius
Mean_Radius

**Supplemental Figure 6. Effect of nutrients on chromatin features and their entropy in protoplast replicate cultures**

**(A)** Principal Component Analysis (PCA) computed on (i) intensity features, (ii) morphology features, (iii) texture features for H2B-RFP during protoplast culturing in nutrient rich (Gamborg) or nutrient poor (W5), each with hormones (+H) or without (-H). The cultures stem from the same, original pool of protoplast partitioned in the different media and imaged at day 0 and day 2 in two or three replicate wells (number of datapoints with the same colour); dataset HTI005. **(B)** PCA computed on all features during protoplast culturing in Gamborg and W5. The cultures stem from the same, original pool of protoplast partitioned in the different media and imaged at day 0, day 3, day 5 and day 7 in at least three replicate wells (number of datapoint with the same colour); right panel: contribution of the families of features to each principal components; dataset HTI001; Right panel: Contribution of the families of features to each principal component. **(C)** H1.2-GFP/H2B-RFP intensity mean ratio measured for protoplasts derived from two different Arabidopsis lines expressing H1.2-GFP/H2B-RFP. **(D)** Effect of the Gamborg and W5 media on nuclei size and shape; P-values Kruskal-Wallis test – all morphology features; left panel: heatmap showing the P value between day 7 and day 0 for each morphology features and pointing out the features with more drastic changes. Right panel: Violin plots for area and roundness. **(E)** Representative images of protoplast nuclei in two different media, sampled at day 7 of culturing, whole ROI and representative nuclei below. **(F)** Differential entropy $\Delta H_0$ between day 7 and day0 for the set of intensity features, morphology features or LBP texture features, respectively, comparing protoplast cultured in the nutrient-rich Gamborg medium or nutrient-poor W5 basal medium. Ns, no significant difference (paired t-test). **(G)** Heatmap showing the P-value (paired t-test) comparing the differential entropy for each single morphology feature and showing a significant effect of nutrients on selective features only (red, orange values)

**A PCA (all features),** two replicate experiments

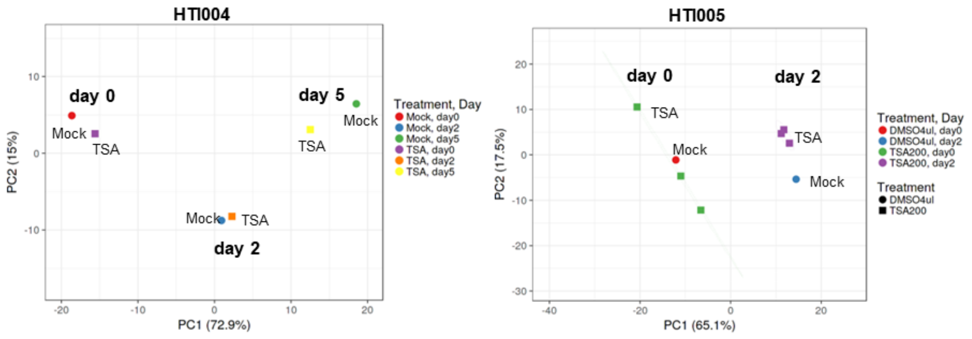


**B H1.2-GFP / H2B-RFP intensity ratio** (HTI004)

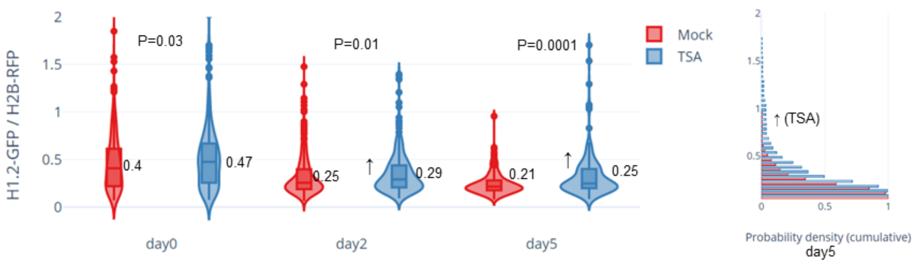


**C Effect of TSA on nuclei size and shape** (HTI004)

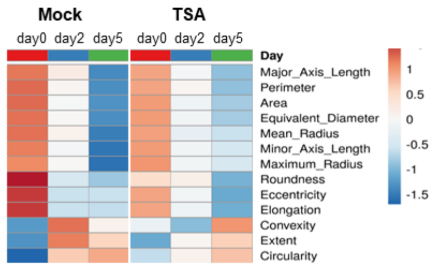


**P-values Kruskal Wallis Mock vs TSA**

| | day0 | day2 | day5 |
|---|---|---|---|
| Area | 0.46 | 0.39 | <u>0.03</u> |
| Circularity | 0.24 | <u>0.01</u> | 0.75 |
| Convexity | 0.25 | 0.01 | 0.09 |
| Eccentricity | 0.20 | 0.78 | 0.73 |
| Elongation | 0.20 | 0.78 | 0.73 |
| Equivalent Diameter | 0.46 | 0.39 | 0.03 |
| Extent | 0.87 | 0.04 | 0.57 |
| Major Axis Length | 0.41 | 0.33 | 0.03 |
| Maximum Radius | 0.63 | 0.15 | <u>0.01</u> |
| Mean Radius | 0.49 | 0.10 | <u>0.02</u> |
| Minor Axis Length | 0.73 | 0.49 | 0.03 |
| Perimeter | 0.38 | 0.94 | <u>0.04</u> |
| Roundness | 0.11 | <u>0.03</u> | 0.88 |

**number of nuclei**

| | day0 | day2 | day5 |
|---|---|---|---|
| **Mock** | 354 | 388 | 217 |
| **TSA** | 327 | 332 | 176 |

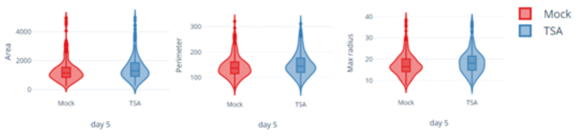


**Supplemental Figure 7. Effect of the histone deacetylase inhibitor TSA on chromatin features and their entropy in protoplast cultures**

**(A)** Principal Component Analysis (PCA) computed on all features for two datasets HTI004 and HTI005; PC1, PC2 - two major principal components; TSA, Trichostatin A (100 nM); Mock – 2% DMSO which was the solvent for TSA; TSA200 – double concentration of TSA (200 nM); Mock – 4% DMSO – double concentration of DMSO. **(B)** Left panel: violin plots for H1.2-GFP:H2B-RFP intensity ratio during protoplast culturing for TSA treatment; right panel: density plot for H1.2-GFP:H2B-RFP intensity ratio at day 5 for TSA treatment. **(C)** Effect of TSA on nuclei size and shape. Left panel: heatmap representing the median value for selected shape descriptors for protoplasts during culturing, treated with TSA and Mock; right panel: p-values calculated with Kruskal Wallis test for selected shape descriptors.

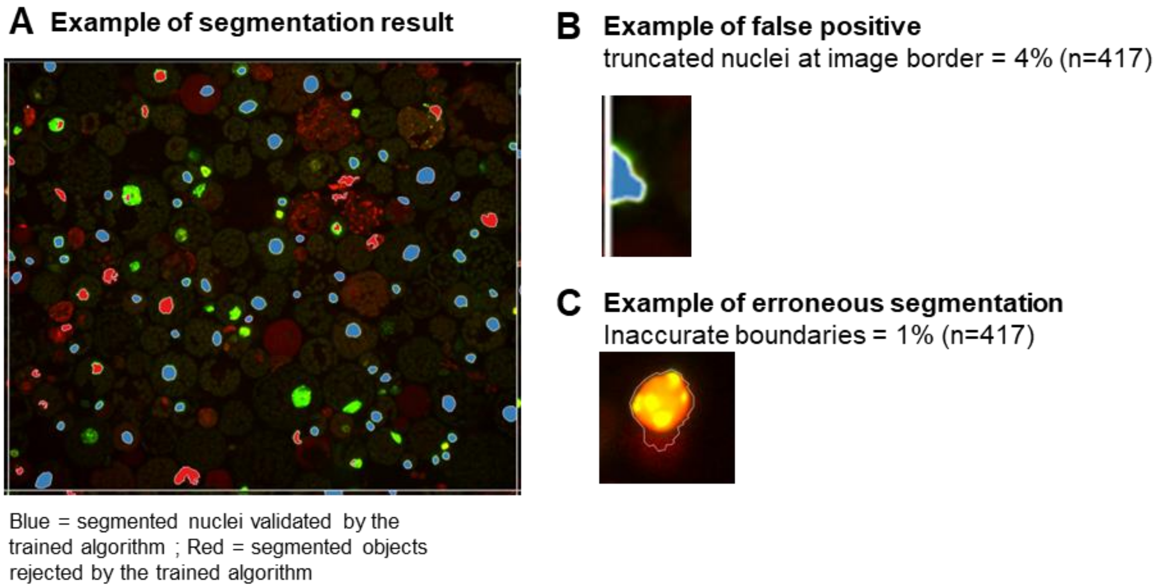


**Supplemental Figure 8. Representative example of nuclei segmentation following training, based on the H2B-RFP signal**

**(A)** Example of segmentation result; blue = segmented nuclei validated by the trained algorithm; red = segmented objects rejected by the trained algorithm. **(B)** Example of the false positive result of the object classification. **(C)** Example of erroneous segmentation.

**Table S2 - Summary of image features exported**

| **Morphology (15)** | **Intensity (5 per channel + 2)** | **Texture (4 methods, 172 metrics per channel)** | | | |
|---|---|---|---|---|---|
| Area | Intensity_max | **_LBP method (108)** | **_TAS (54)** | **_Hu (7)** | **_Gabor method (3)** |
| Circularity | Intensity_mean | radius 1-10.9 in steps of 0.01 to 0.1 | center, mu, mu.margi, n.center, n.mu, n.mumargin radius from 0 to 8 | steps 0 to 6 | frequency 1, 5, 10 |
| Convexity | Intensity_min | | | | |
| Eccentricity | Intensity_sum | | | | |
| Elongation | Intensity_std | | | | |
| EquivalentDiameter | Channel Ratio_Mean_Intensity | | | | |
| Extent | Channel Ratio_Sum_Intensity | | | | |
| LocalCentroidx | | | | | |
| LocalCentroidy | | | | | |
| MajorAxisLength | | | | | |
| MinorAxisLength | | | | | |
| MaximumRadius | | | | | |
| MeanRadius | | | | | |
| Perimeter | | | | | |
| Roundness | | | | | |

**Table S3**. Media composition.

Gamborg B5 Basal - Contains the macro- and micronutrients, and vitamins as described by Gamborg, et al. (1968).

| | **Basic media** | | Modified media | | | |
|---|---|---|---|---|---|---|
| | **W5** | **Gamborg** | W5 + hormones | Gamborg without hormones | Gamborg2 | Gamborg2 without hormones |
| **Reagents** | [ml] | [ml] or [g] | [ml] | [ml] or [g] | [ml] or [g] | [ml] or [g] |
| *Total volume* | *50* | *50* | *50* | *50* | *50* | *50* |
| 5M NaCl | 1.54 | | 1.54 | | | |
| 2M KCl | 0.125 | | 0.125 | | | |
| 1M $CaCl_2$ | 6.25 | 0.34 | 6.25 | 0.34 | 0.147 | 0.147 |
| 0.2M MES | 0.5 | 0.8 | 0.5 | 0.8 | 0.8 | 0.8 |
| Ampicilin | 0.05 | 0.05 | 0.05 | 0.05 | 0.05 | 0.05 |
| 2,4D (1mg/ml) | | 0.05 | 0.05 | 0.05 | 0.05 | |
| BAP (1.5mg/ml) | | 0.005 | 0.005 | 0.005 | | |
| Kinetin (1 mg/ml) | | | | | 0.005 | |
| Gamborg B5 Basal | | 0.16 g | | 0.16 g | 0.16 g | 0.16 g |
| Glucose | | 3.6 g | | 3.6 g | 2 g | 2 g |
| Mannitol | | | | | 3 g | 3 g |

**Table S4 - Datasets used for the study**

| Dataset Name (csv file) | Internal Name | Time Points | Imaging days | Cultures (wells) | Medium | Treatment | Line | Line_description | Image file |
|---|---|---|---|---|---|---|---|---|---|
| HTI_001.csv | exp4 | 4 | day0, day3, day5, day7 | B02-B05 | Gamborg | none | KR239 | H1.2-GFP/UBQ10::H2B-RFP | IDR001.zip |
| HTI_001.csv | | | | C02-C05 | W5 | none | KR239 | H1.2-GFP/UBQ10::H2B-RFP | IDR001.zip |
| HTI_001.csv | | | | D02-D05 | Gamborg | none | KR236 | H1.2-GFP/UBQ10::H2B-RFP | IDR001.zip |
| HTI_001.csv | | | | E02-E05 | W5 | none | KR236 | H1.2-GFP/UBQ10::H2B-RFP | IDR001.zip |
| HTI_002.csv | exp5 | 3 | day0, day2, day4 | B06-B07 | Gamborg | none | KR318 | HTR5::YFPnls/UB10::H2B-RFP | IDR002.zip |
| HTI_002.csv | | | | B09-B10 | Gamborg | none | KR317 | HTR5::YFPnls/UB10::H2B-RFP | IDR002.zip |
| HTI_002.csv | | | | C05-C07 | W5 | none | KR317 | HTR5::YFPnls/UB10::H2B-RFP | IDR002.zip |
| HTI_002.csv | | | | C08-C10 | W5 | none | KR318 | HTR5::YFPnls/UB10::H2B-RFP | IDR002.zip |
| HTI_004.csv | exp7a | 3 | day0, day2, day5 | B2-B4 | Gamborg | none | KR239 | H1.2-GFP/UBQ10::H2B-RFP | IDR004.zip |
| HTI_004.csv | | | | B5 | Gamborg | FDA at day0 | KR239 | H1.2-GFP/UBQ10::H2B-RFP | IDR004.zip |
| HTI_004.csv | | | | B6 | Gamborg | FDA at day2 | KR239 | H1.2-GFP/UBQ10::H2B-RFP | IDR004.zip |
| HTI_004.csv | | | | B7 | Gamborg | FDA at day5 | KR239 | H1.2-GFP/UBQ10::H2B-RFP | IDR004.zip |
| HTI_004.csv | | | | B8-B10 | Gamborg | TSA 100nm | KR239 | H1.2-GFP/UBQ10::H2B-RFP | IDR004.zip |
| HTI_004.csv | | | | B11 | Gamborg | DMSO 2% | KR239 | H1.2-GFP/UBQ10::H2B-RFP | IDR004.zip |
| HTI_004.csv | | | | C2-C4 | Gamborg/noHormones | none | KR239 | H1.2-GFP/UBQ10::H2B-RFP | IDR004.zip |
| HTI_005.csv | exp7b | 2 | day0, day2 | D2-D4 | W5/with_hormones | none | KR239 | H1.2-GFP/UBQ10::H2B-RFP | IDR005.zip |
| HTI_005.csv | | | | D8 | Gamborg | TSA 100 nM at day1 | KR239 | H1.2-GFP/UBQ10::H2B-RFP | IDR005.zip |
| HTI_005.csv | | | | D9-D10 | Gamborg | none | KR239 | H1.2-GFP/UBQ10::H2B-RFP | IDR005.zip |
| HTI_005.csv | | | | D11 | Gamborg | DMSO 2% at day1 | KR239 | H1.2-GFP/UBQ10::H2B-RFP | IDR005.zip |
| HTI_005.csv | | | | E4 | W5 | none | KR239 | H1.2-GFP/UBQ10::H2B-RFP | IDR005.zip |
| HTI_005.csv | | | | E5 | W5 | none | KR239 | H1.2-GFP/UBQ10::H2B-RFP | IDR005.zip |
| HTI_005.csv | | | | E9-E10 | Gamborg | TSA 200nM | KR239 | H1.2-GFP/UBQ10::H2B-RFP | IDR005.zip |
| HTI_005.csv | | | | E11 | Gamborg | DMSO 4% | KR239 | H1.2-GFP/UBQ10::H2B-RFP | IDR005.zip |

### Supplementary Information File 1 – Entropy Script

```
# Compute entropy by subsampling

########################################################
########################################################
#Estimate of the differential entropy
ent_bin= function(Y,indices=NULL,larg_bin=NULL){
  #Compute the h_corrected estimator
  #if larg_bin = NULL -> use the Freedman-Diaconis method to construct bins
  #can be used for bootstrap sampling with the library 'boot'
  if (is.null(indices)){indices = 1:length(Y)}
  X = Y[indices]
  if (is.null(larg_bin)){
    if (IQR(X) == 0){stop('methode de decoupage FD inutilisable')}
    larg_bin = 2*IQR(X)/(length(X)^(1/3))
  }
  bin = floor((X-min(X))/larg_bin)+1
  p = (1/length(X))*table(bin)
  id = (p>0)
  H_discr=-sum(p[id]*log(p[id]))+log(larg_bin)
  return(H_discr)
}
########################################################
########################################################



########################################################
########################################################
#BUB estimator for discrete entropy
#Mathilde GAILLARD
#Ref. Liam PANINSKI Estimation of Entropy and Mutual Information
#
library(pracma)
library(corpcor)
bub_opti = function(N,m,k_max){
  #m = nber of bins
  #N = nber of data
  #k_max = nber of coefficients a_j for which bub_opti is using the BUB method otherwise
-> compute MM estimator
  #k_max should be less ofr equal to ? N

  p = (0:N)/N
  B = mat_bernouilli_polynom(p,N)
  g = sapply(p,f,m)
  g_col = matrix(g, ncol = 1)
  h = c(0, sapply(p[-1],H))
  Y_tot = matrix(h, ncol = 1)

  #every a_j initialized with the MM estimator
  a = (0:N)/N
  a = -a*log(a)+((1-a)/(2*N))

  best_MM = Inf
  for (i in (1:k_max)){
    #print(i)
    h_exp = a[(i+1):length(a)]%*%(B[(i+1):length(B[,1]),])#part of the entropy already
explained by the coeff
    Y_non_pond = Y_tot-t(h_exp) #withdraw the part explained

    G = repmat(g,i,1)
    X = t(G*B[1:i,])
    D = Diag(rep(-1,i),0)+Diag(rep(1,i-1),1)
    U = (t(X)%*%X)+(N/4)*(t(D)%*%D)
    U[i,i] = U[i,i]+(N/4)

    Y = g_col*Y_non_pond
    XY = (t(X)%*%Y)#-h_exp
    XY[i] = XY[i] + (N/4)*a[i+1]

    a[1:i] = pseudoinverse(U)%*%XY

    #make a 1 col matrix
    a_mat_1col = matrix(a, ncol = 1)

    #compare the performance of this specific set f a_j
    biais = 2*g*abs((t(Y_tot)-a%*%B))
```

```
    maxbiais = max(abs(biais))
    borne_var = max((a-c(a[2:length(a)],0))^2)
    MM = sqrt(maxbiais^2+N*borne_var)

    if(MM<best_MM){
      #choose the best set of a_j
      best_MM = MM
      best_a = a
      best_biais = maxbiais
    }
  }
  return(best_a)
}

mat_bernouilli_polynom = function(p,N){
  #compute the matrix of bernouilli polynomials
  fa = lgamma(1:(N+1))
  Ni = fa[N+1]-fa[1:(N+1)]-rev(fa)

  B = zeros(n =  N+1, m = N+1)
  p = p[-1]
  p = p[-length(p)]

  lp = log(p)
  lq = rev(lp)

  for (i in 0:N){
    B[i+1,2:N] = Ni[i+1]+i*lp+(N-i)*lq
  }
  B = exp (B)
  B[2:(N+1),1]=zeros(N,1)
  B[1:N,N+1]=zeros(N,1)
  return (B)
}

H = function(x){
  #entropy function
  nz = (x>0)
  return (-sum(x[nz]*log(x[nz])))
}

f = function(x,m){
  #weight function
  if(x<(1/m)){
    return(m)
  }
  else{
    return(1/x)
  }
}

estimateur_bub_entropie_discrete = function(X){
  #compute the discrete entropy with the bub_opti function
  N = length(X)
  h = tabulate (X+1)
  m = length(h)
  hist_eff = c()
  for (i in 1:(N+1)){
    hist_eff[i]=sum((h==i-1))
  }
  A = bub_opti(N,m,k_max = 11) #k_max = 11 works when N<10^6
  H_BUB = sum(A*hist_eff)
  return(H_BUB)
}
#########################################################
#########################################################



#########################################################
#########################################################
# Define subsampling for discrete entropy
subsampled_BUB_H = function(X,nr,msd){
  #compute the discrete entropy with the bub_opti function
  H0={}
  for (i in 1:nr) {
    s1=sample(X,msd,replace=F)
    E=estimateur_bub_entropie_discrete(s1)
    H0=append(H0,E)
```

```
    print(nr-i)
  }
  return(mean(H0))
}
#########################################################
#########################################################


#########################################################
#########################################################
# Define subsampling for differential entropy
subsampled_bin_H = function(X,nr,msd){
  #compute the differential entropy
  H0={}
  for (i in 1:nr) {
    s1=sample(X,msd,replace=F)
    E=ent_bin(s1)
    H0=append(H0,E)
    print(nr-i)
  }
  return(mean(H0))
}
#########################################################
#########################################################

# Read the file
path="/PATH/DATASET.csv"
setwd(path)
dr=read.table("DATASET.csv",header=T, sep=';')

# Determine the smallest time point in cell number
sd0=sum(dr$Day=="day0")
sd2=sum(dr$Day=="day2")
sd5=sum(dr$Day=="day5")
msd=min(sd0,sd2,sd5)

Day=c("day0", "day2", "day5")

# test mode
nr=4
nc=4

# production mode
nc=ncol(dr)
nr=100

H_0={}
H_1={}
for (i in 24:nc) {
  if (typeof(dr[,i]) == "integer") {
    print ("I")
    for (j in 1:3) {
      M=(dr[,i][dr$Day==Day[j]])
      E=subsampled_BUB_H(M,nr,msd)
      H_0=append(H_0, mean(E))
    }
    print (H_0)
  }
  else {
    print ("D")
    for (j in 1:3) {
      M=(dr[,i][dr$Day==Day[j]])
      if(IQR(M)==0) {
        i=i+1
        H_0=H_0=append(H_0, "NA")
      }
      else {
        E=subsampled_bin_H(M,nr,msd)
        H_0=append(H_0, mean(E))
      }
    }
    print (H_0)
  }
  H_1=cbind(H_1,H_0)
  H_0={}
}

colnames(H_1)=colnames(dr)[4:ncol(dr)]
write.table(H_1, "entropy_values")
```
